# Persistent antigen is essential for sustaining *Leishmania*-specific memory CD4^+^ T cells and long-term immunity

**DOI:** 10.1101/2025.01.26.633721

**Authors:** Zhirong Mou, Roma Zayats, Enitan Salako, Nnamdi Ikeogu, Somtochukwu S Onwah, Dong Liu, Hiroshi Hamana, Da Tan, Hiroyuki Kishi, Carson Leung, Janilyn Arsenio, Thomas Murooka, Jude E Uzonna

## Abstract

Memory T cells are critical for secondary immunity against pathogens, yet their persistence in the absence of antigen remains unclear. While memory CD8^+^ T cells are known to persist independently of their cognate antigen, the durability of memory CD4^+^ T cells in the absence of antigen remains controversial. Recovery from cutaneous leishmaniasis confers lifelong immunity, largely mediated by CD4^+^ T cells, but the necessity of persistent parasites for sustaining this immunity has not been empirically confirmed. We investigated immunity in mice infected with a dihydrofolate reductase-thymidylate synthase (*dhfr-ts*)-deficient *Leishmania major*, which cannot persist due to their thymidine salvage deficiency. These mice lost protection against wild-type challenge, correlating with a decline in *Leishmania* (PEPCK)-specific CD4^+^ T cells. To further dissect this relationship, we generated PEPCK-specific CD4^+^ TCR transgenic (PEG) mice, enabling precise tracking of *Leishmania*-specific memory CD4^+^ T cells. Both *in vitro* and *in vivo*-generated memory PEG cells gradually disappeared over time in the absence of antigen, irrespective of the host’s MHC II status, and this loss paralleled the erosion of infection-induced immunity. PEG and endogenous PEPCK-specific memory cells were not maintained in mice infected with *dhfr-ts*- or *PEPCK*-deficient *L. major*, resulting in loss of recall responses and secondary immunity. These findings demonstrate that continuous antigen presence is crucial for maintaining *Leishmania*-specific memory CD4^+^ T cells and highlight the role of persistent antigen in sustaining long-term immunity against chronic infections.

## INTRODUCTION

Immunologic memory is a hallmark of adaptive immunity and the basis for vaccine design and vaccination strategies against infectious agents. It ensures an accelerated and enhanced immune response upon re-infection with the same pathogen. Memory T cells play essential roles in immunity against virus, intracellular bacteria, and protozoan parasites. Three traditional features of memory T cells include prior activation and expansion, persistence in the absence of cognate antigen, and enhanced functional activity upon antigen re-exposure ^1^. However, not all of these features are well validated due to lack of precise definitions of memory cells and their functional or phenotypic characteristics ^2^, especially for CD4^+^ memory T cells. Although CD8^+^ T cells are maintained as a stable memory pool for extended time periods ^1,3,4^, whether memory CD4^+^ T cells are sustained in the absence of their cognate antigens is controversial. While some studies show that the numbers of memory CD4^+^ T cells decline over time during LCMV and *Listeria* infections ^4,5^, others found that memory CD4^+^ T cells numbers remain stable over time ^6^. Also, BCG, which is the best available tuberculosis vaccine, does not provide children with life-long CD4^+^ T cell immunity and is not effective in adults ^7^, and mumps outbreaks occur in vaccinated individuals ^8^. Furthermore, while acquired immunity to malaria is maintained in adults living in endemic areas, those who left endemic regions and migrate to malaria-free countries lose such immunity and become highly susceptible upon return to the endemic areas ^9,10^. These observations question the dogma that maintenance of immunological memory (particularly memory CD4^+^ T cells) is independent of their cognate antigen.

Leishmaniasis is caused by the protozoan parasite belonging to the genus *Leishmania*. Most individuals who recover from cutaneous leishmaniasis, the most common form of the disease, acquire lifelong immunity to reinfection. This infection-induced immunity is mediated by IFN-γ-producing CD4^+^ T cells and dependent on persistence of parasites at primary infection site ^11–13^. The findings that *L. major*-infected IL-10-deficient mice are able to completely clear parasites ^14^ but also lose immunity to reinfection ^15^, suggests that persistent antigens are important for the maintenance of protective *Leishmania*-specific CD4^+^ T cells and hence immunologic memory against *Leishmania* parasites.

Here, we provide compelling data that unequivocally demonstrate that both endogenous and transgenic *Leishmania*-specific memory CD4^+^ T cells do not persist in the absence of their cognate antigen. This loss of *Leishmania*-specific memory CD4^+^ T cells is associated with loss of infection-induced immunity resulting in increased susceptibility to secondary *L. major* challenge. These observations are inconsistent with the dogma that long-term maintenance of antigen-specific memory CD4^+^ T cells is independent of their cognate antigens.

## RESULTS

### *Leishmania*-specific CD4^+^ T cells that mediate infection-induced immunity are lost over time in the absence of *Leishmania* antigen

Humans and mice that recover from cutaneous leishmaniasis acquire lifelong immunity to reinfection and this is associated with persistence of few parasites at the primary infection site. This memory CD4^+^ T cell-mediated infection-induced immunity is lost following complete clearance of parasites ^13,14^, suggesting that persisting parasites (antigen) may be critical for their maintenance. To unequivocally determine if clearance of parasites (antigen) is responsible for this loss of protection, we used *dhfr-ts^-^ L. major* parasites, which elicit strong early immunity, but are unable to persist for a long time in infected mice ^16^. Mice infected with *dhfr-ts^-^ L. major* for 6 weeks displayed similar protection as those infected with wild-type (WT) parasites against secondary virulent *L. major* challenge as evidenced by strong DTH responses and lower parasite burden at the challenge site (Fig. 1A-C). However, after 24 weeks (when *dhfr-ts^-^* parasites are completely cleared ^16,17^), there was loss of protection (DTH response and parasite control, Fig. 1D and 1E) against secondary virulent challenge in mice infected with *dhfr-ts^-^ L. major*, but not in those infected with WT parasites or *dhfr-ts^-^* spiked with WT parasites. These observations suggest that persistent parasites may play a crucial role in maintenance of *Leishmania*-specific memory CD4^+^ cells that mediate secondary protection against *L. major* challenge. Infection-induced immunity can be adoptively transferred to naïve animals by transferring splenocytes or CD4^+^ T cells from *L. major*-infected and subsequently healed animals ^17,18^. To further determine the importance of persistent parasites (antigen) in maintenance of infection-induced immunity, we adoptively transferred splenocytes from *L. major*-infected and healed mice into naïve mice and challenged them with virulent *L. major* at 1, 7, or 24 weeks post cell transfer (Fig. 2A). Recipient mice displayed strong DTH responses (Fig. 2B) and lower parasite burden (Fig. 2C) when challenged at 1 and 7 weeks post cell transfer. In contrast, DTH response or protection were lost when recipient mice were challenged at 24 weeks post cell transfer (Fig. 2C), indicating that the protective *Leishmania*-specific CD4^+^ T cells did not persist for a long time in the recipient animals. In line with this, *Leishmania* PEPCK-specific CD4^+^ T cells, which are dominant and protective against *L. major* infection in mice ^18^, were detectable at 1 week in the recipient animals (Fig. 2E), but not at 24 weeks (Fig. 2E and 2F) after cell transfer. Collectively, these results strongly indicate that maintenance of *Leishmania* antigen-specific memory CD4^+^ T cells that mediate durable infection-induced immunity is dependent on the presence of their cognate antigens.

**Fig. 1.**
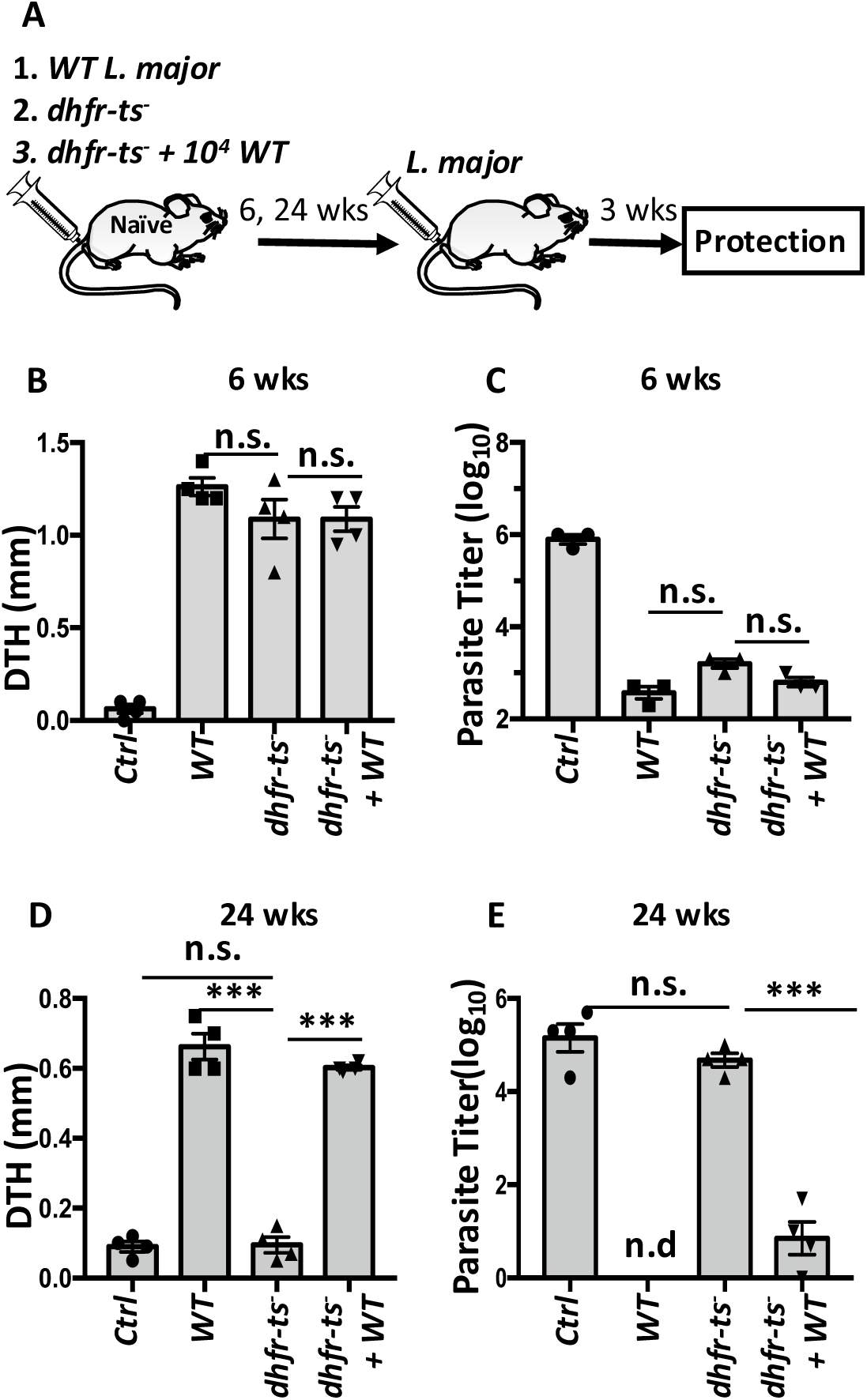
Infection-induced immunity mediated by auxotrophic *dhfr-ts^-^ L. major* is lost following parasite clearance. (**A**) Schematic protocol of the experimental design. Eight weeks old C57BL/6 mice were infected with 10^6^ WT, *dhfr-ts^-^* or 10^6^ *dhfr-ts^-^* spiked with 10^4^ WT *L. major* in the right hind footpads. At 6 (**B-C**), or 24 (**D-E**) weeks post-infection, the mice were rechallenged in the left hind footpads with 10^6^ WT *L. major*. Footpad swelling (DTH response) was measured at 3 days post-rechallenge (**B, D**) and mice were sacrificed at 3 weeks to quantify parasite burden (**C, E**). Results presented are representative of 2-3 independent experiments (n = 4 mice in each group per experiment) with similar results. *, p < 0.05; **, p < 0.01; ***, p < 0.001; n.s., not significant.

**Figure 2.**
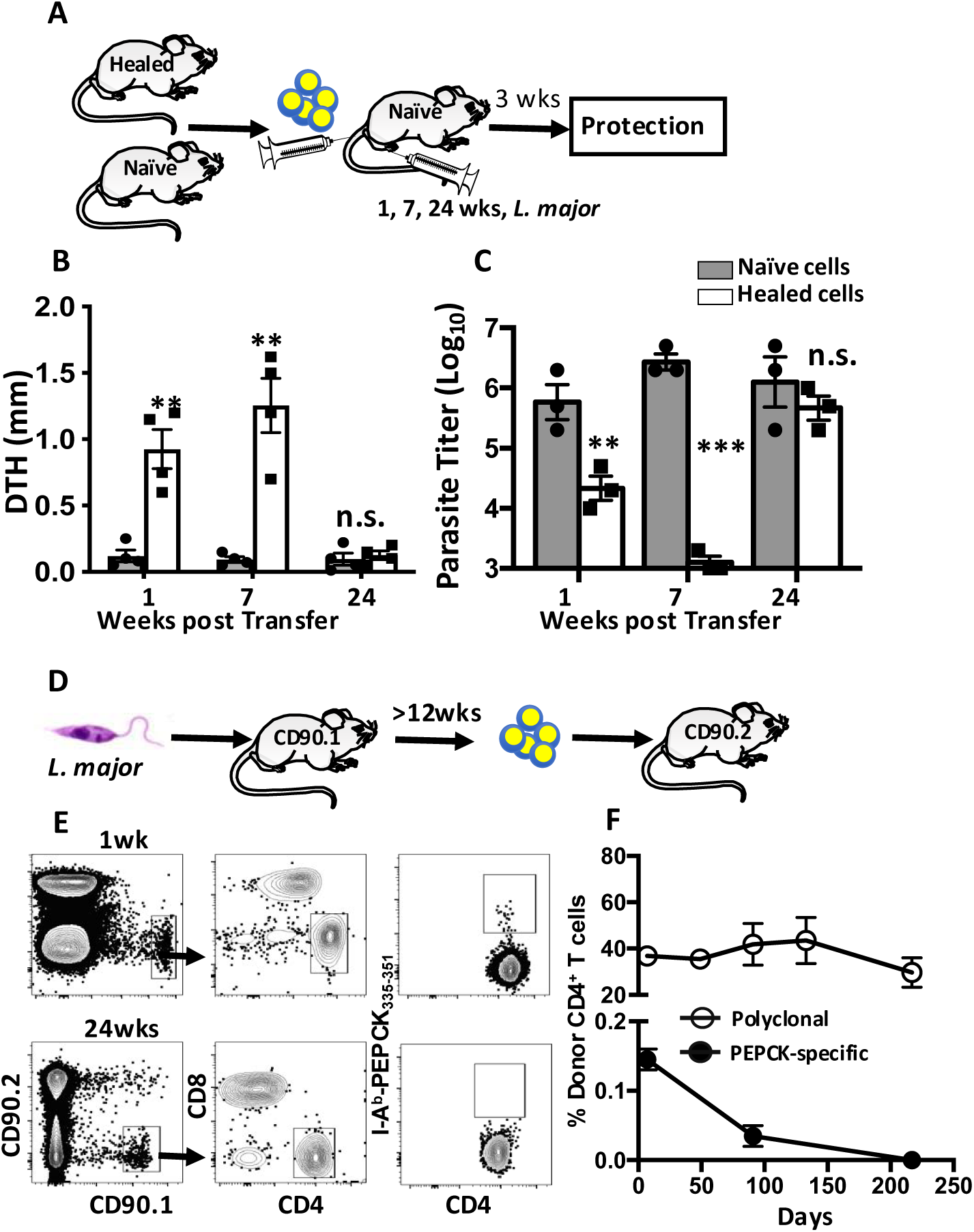
Adoptively transferred infection-induced immunity is lost following loss of *Leishmania*-specific (PEPCK-specific) CD4^+^ T cells in the recipient mice. (**A**) Schematic protocol showing experimental design used to assess adoptive transfer of infection-induced immunity to naïve recipient mice. Five million (5 ξ 10^6^) splenic CD4^+^ T cells from naïve or healed (previously infected with WT *L. major* for more than 12 weeks) mice were injected intravenously into naïve C57BL/6 (recipient) mice. Recipient mice were challenged with WT *L. major* at 1, 7 and 24 weeks post-cell transfer and DTH response (**B**) was measured after 3 days. Three weeks after challenge, mice were sacrificed and parasite burden (**C**) in the challenged footpads was measured. To monitor longevity of PEPCK specific-CD4^+^ T cells, 2 ξ 10^7^ splenocytes from infected and healed CD90.1^+^ C57BL/6 mice were adoptively transferred into naïve congenic (CD90.2^+^) mice (**D**). Total donor CD4^+^ or PEPCK specific-CD4^+^ T cells were determined at different times post transfer by flow cytometry (**E-F**). Results presented are representative of 2-3 independent experiments (n = 3-4 mice in each group per experiment) with similar results. **, p < 0.01; ***, p < 0.001; n.s., not significant.

### Generation and characterization of PEG mice

The above results strongly suggest that antigen persistence is required for maintenance of *Leishmania*-responsive memory CD4^+^ T cells. To unequivocally demonstrate this, we generated *L. major* PEPCK-specific T cell receptor (TCR) transgenic (Tg) mice (PEG) following our identification of PEPCK as an immunodominant antigen of *Leishmania* ^18^. We cloned PEPCK-specific CD4^+^ TCR alpha and beta genes by single-cell PCR technique ^18^, transduced several clones into TG 40 cell lines (Fig. S1A and B) or primary CD4^+^ T cells (Fig. S1C-1E) and demonstrated that they bind to I-A^b^-PEPCK_335-351_ tetramer (Fig. S1B and 1C), proliferate (Fig. S1D) and produce IFN-γ (Fig. S1E) in response to PEPCK_335-351_ peptide. TCR31 clone was selected and subcloned into VA hCD2 vector ^19^ (Fig. 3A and Fig. S2A) to generate TCR Tg mice (PEG) by pronuclear injection. Over 90% of CD4^+^ T cells in PEG mice express the transgenic TCR as evidenced by reactivity with I-A^b^-PEPCK_335-351_ tetramer (Fig. S2B). PEG mice had slightly more CD4^+^ T cells (Fig. S2C) and fewer natural Treg cells (Fig. S2D) than their WT mice. PEG CD4^+^ T cells also differentiated into Th1, Th17 and Treg cells *in vitro* akin to their WT counterpart mice (Fig. S2E).

**Figure 3.**
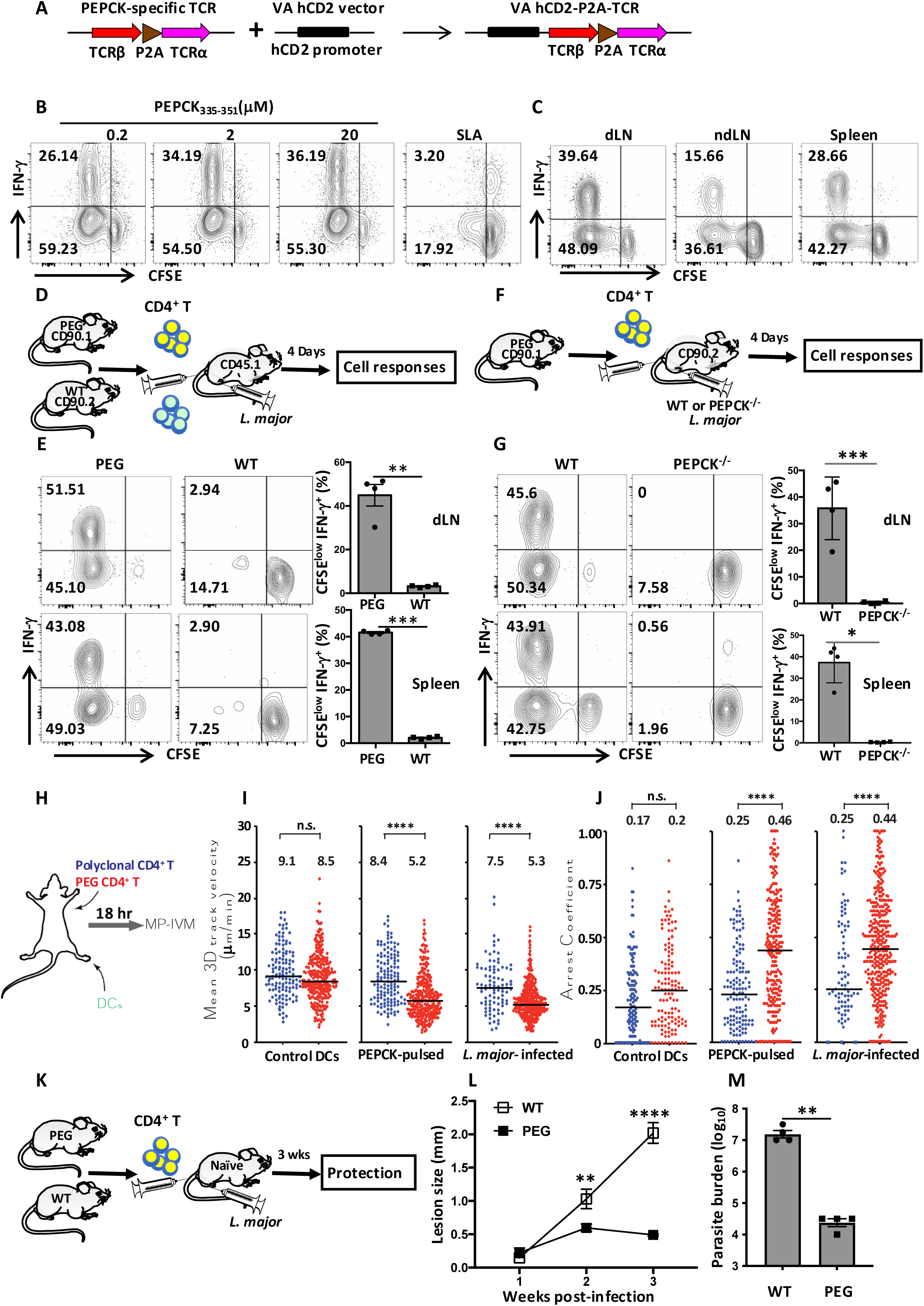
Generation and characterization of PEG mice. (**A**) Schematic protocol for generating PEPCK-specific TCR transgenic (PEG) mice. PEPCK-specific TCR alpha and beta genes linked with P2A linker were subcloned into VA hCD2 vector under hCD2 promoter. Linearized transgenic plasmid DNA fragments were inoculated (by pronuclear injection) into fertilized C57BL/6 eggs that were then implanted into surrogate mothers to obtain founder offspring. (**B**) CD4^+^ T cells from PEG mice proliferated and produced IFN-γ in response to PEPCK_335-351_ *in vitro*. CD4^+^ T cells from PEG mice were labelled with CSFE dye and co-cultured with BMDCs in the presence of different concentrations of PEPCK_335-351_ peptide. After 5 days, proliferation and IFN-γ production were measured by flow cytometry. (**C**) PEG cells respond *L. major* expressing PEPCK antigen *in vivo*. One million (10^6^) purified CFSE-labelled PEG CD4^+^ T cells were adoptively transferred into congenic mice. On next day, the recipient mice were challenged with 1 x 10^6^ *L. major* in the footpads and proliferation and IFN-γ production were measured after 4 days. (**D** and **E**) PEG CD4^+^ T cells are functional *in vivo*. CFSE-labelled naïve PEG (CD90.1^+^CD45.2^+^) and polyclonal WT (CD90.2^+^ CD45.2^+^) CD4^+^ T cells were mixed in equal proportion and co-transferred into congenic CD45.1^+^ recipients that were challenged with *L. major* the next day. After 4 days, mice were sacrificed, and proliferation and IFN-γ production were measured. (**F** and **G**) PEG CD4^+^ T cells are PEPCK-specific. CFSE-labelled naïve PEG CD4^+^ T cells were transferred into recipient mice, challenged with either WT or PEPCK deficient (PEPCK^-/-^) *L. major* and sacrificed after 4 days to assess proliferation and IFN-γ production. (**H-J**) MP-IVM analysis of DC:T cell dynamics in the popliteal lymph node. Dendritic cells (DCs) were pulsed with PEPCK_335-351_ peptides or infected with *L. major* and injected into the footpads of naïve recipient animals. After 12 hours, the mice were injected with either naïve polyclonal or PEG CD4^+^ T cells that were labeled with different Celltracker dyes. Cell interactions were imaged in the popliteal lymph node next day by intravital microscopy. (**H**) Mean 3D track velocity (**I**) and arrest coefficient (**J**) of polyclonal (blue) and PEG (red) CD4^+^ T cells compared to control, PEPCK_335-351_ pulsed or *L. major*-infected BMDCs were analyzed. Indicated numbers and black lines are median values. Data are pooled from 11 individual recordings from 6 animals. (**K-M**) PEG CD4^+^ T cells protect mice against virulent *L. major* challenge. One million CD4^+^ T cells from naïve WT (polyclonal) or PEG mice were adoptively transferred into naïve C57BL/6 mice that were then challenged with 10^6^ virulent *L. major* (**K**). Lesion size was monitored weekly (**L**) and mice were sacrificed after 3 weeks to determine parasite burden was measured by limited dilution (**M**). Results presented are representative of 2-3 independent experiments (n = 4-6 mice in each group per experiment) with similar results. *, p < 0.05; **, p < 0.01; ****, p < 0.0001. n.s., not significant.

PEG CD4^+^ T cells proliferated and produced IFN-γ in response to PEPCK_335-351_ peptide presentation by DCs *in vitro* (Fig. 3B) and *L. major* infection *in vivo* (Fig. 3C). To determine if PEG CD4^+^ T cell responses are specific, PEG cells were transferred either alone or mixed with WT CD4^+^ T cells into congenic mice (Fig. 3D and 3E) and challenged with WT or PEPCK deficient (PEPCK^-/-^) *L. major*. PEG (but not WT), CD4^+^ T cells proliferated and produced IFN-γ (Fig. 3E) in response to WT *L. major* challenge. In contrast, PEG CD4^+^ T cells did not respond to PEPCK^-/-^ *L. major* challenge (Fig. 3F and 3G), confirming their antigen specificity.

Next, we recorded the migratory behavior of PEG CD4^+^ T cells in the draining popliteal lymph node by multiphoton intravital microscopy (MP-IVM) and found that they displayed reduced 3D migration speeds and higher arrest coefficient in the presence of PEPCK-pulsed or *L. major*-infected DCs compared to control WT T cells (Fig. 3H-3J, Fig. S3). Prolonged DC:T cell clusters are observed in the presence of cognate antigen 18 hours post PEG T cell transfer, indicating antigen-driven stable contacts that are required for T cell activation and expansion ^20^ (Movie S1). Indeed, adoptive transfer of PEG cells into naïve mice strongly protected recipient mice against *L. major* challenge (Fig. 3K) as evidenced by significantly smaller lesion size (Fig. 3L) and lower parasite burden (Fig. 3M). As expected, PEG mice were highly resistant to primary (Fig. S4A and 4B) and secondary *L. major* infection (Fig. S4C). This resistance was associated with high IFN-γ production in the dLN (Fig. S4D) and spleens (Fig. S4E). Collectively, these results demonstrate that PEG CD4^+^ T cells are specific to *L. major* PEPCK and mediate protection against *L. major* challenge akin to PEPCK-specific CD4^+^ T cells generated in *L. major-* infected mice ^18^.

### *Leishmania*-specific (PEG) memory CD4^+^ T cells generated *in vitro* do not persist in wild type (WT) and MHC II deficient (MHC II^-/-^) mice

To address if *Leishmania*-specific memory CD4^+^ T cells can be maintained in the absence of antigen, we generated memory CD4^+^ T cells *in vitro* as previously described ^6^ with slight modification (Fig. 4A, see materials and methods) and transferred them into MHC II^-/-^ and WT (MHC II sufficient) mice. Prior to transfer, *in vitro* generated memory PEG CD4^+^ T cells maintained their reactivity to bind I-A^b^-PEPCK_335-351_ tetramer (Fig. S5A) and expressed markers consistent with effector and central memory cells (Fig. 4B). In addition, *in vitro* generated memory PEG CD4^+^ T cells produce IFN-γ only following restimulation with PEPCK_335-351_ peptide (Fig. 4C) or upon stimulation with PMA and ionomycin (Fig. 4D), indicating that they display characteristics of bona fide memory T cells.

**Fig. 4.**
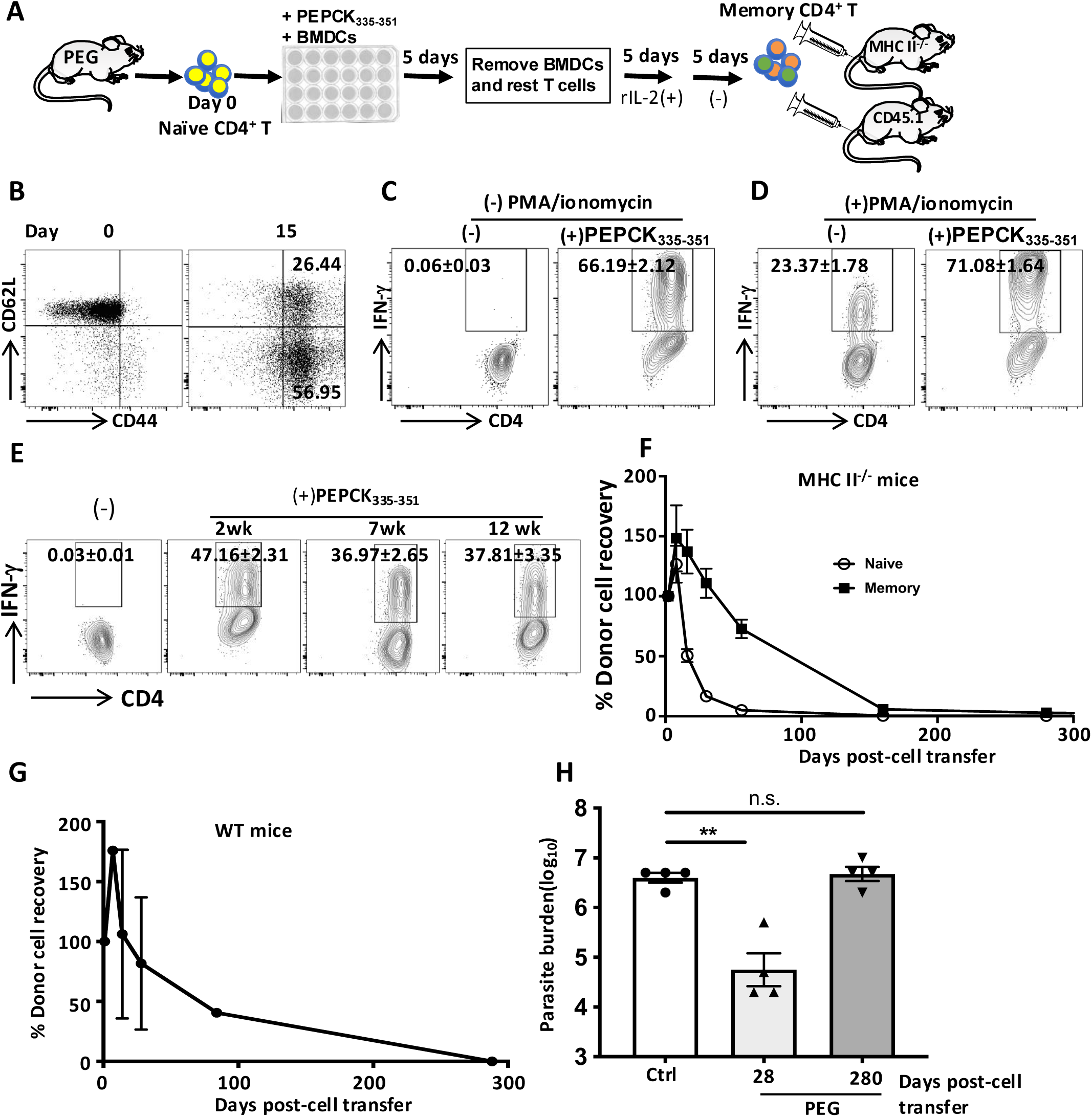
Memory PEG cells generated *in vitro* do not persist in MHC II^-/-^ and WT mice. (**A**) Schematic protocol for generating memory PEG cells *in vitro* adapted from ref 6. (**B**) Expression of CD44 and CD62L on Day 0 (onset of culture) and Day 15 (prior to transfer) of *in vitro* generated memory cells. *In vitro* generated memory PEG cells were stimulated with splenic DCs pulsed with or without PEPCK_335-351_ (5 μM) overnight and then incubated with either BFA alone (**C**) or PMA, ionomycin and BFA (**D**) for additional 4-6 hr and assessed for IFN-γ production by flow cytometry. (**E**) One million memory PEG cells generated *in vitro* were transferred into MHC II^-/-^ mice. At indicted time points, some recipient mice were sacrificed, and PEG cells were reisolated from spleens and lymph nodes, stimulated overnight with WT splenic DCs pulsed with or without PEPCK_335-351_ peptide (5 μM) and incubated with BFA for additional 4-6 hr before assessing for IFN-γ production by flow cytometry. One million naïve or *in vitro* generated memory PEG cells were transferred into MHC II^-/-^ (**F**) or congenic WT (**G**) mice and their survival was monitored over time. At the indicated times, age-matched control naïve mice or WT mice that received *in vitro* generated memory PEG cells were challenged with 10^6^ virulent *L. major* and parasite burden was assessed after 3 weeks post challenge (**H**). Results presented are representative of 2-3 independent experiments (n = 4 mice in each group per experiment) with similar results. *, p < 0.05; n.s., not significant.

Next, we assessed the numbers of *in vitro* generated memory PEG CD4^+^ T cells in MHC II^-/-^ recipient mice. At different time points after cell transfer, memory PEG CD4^+^ T cells recovered from MHC II^-/-^ mice produced IFN-γ in response to PEPCK_335-351_ peptide (Fig. 4E), indicating they were still functional. The number of memory PEG CD4^+^ T cells increased in the first 2 weeks (peaked on 4 days) after cell transfer consistent with homeostatic expansion and reduced to 50% in about a month (Fig. 4F). By 150 days post transfer, 95% and 99.5% of transferred memory and naïve PEG T cells, respectively, have disappeared (Fig. 4F and Fig. S5B-5D). Thus, the numbers of transferred memory PEG cells were not stable over time, which is contrary to a previous report ^6^. Similar results were also observed following transfer of *in vitro* generated memory PEG CD4^+^ T cells into WT mice (Fig. 4G and Fig. S5E and S5F). Interestingly, the disappearance of memory PEG CD4^+^ T cells in the WT recipient mice was associated with loss of protection following virulent *L. major* challenge (Fig. 4H).

To determine if the disappearance of memory cells was a unique characteristic of PEG cells, we generated memory OT II cells (Fig. S6A-6C) and assessed their persistence in MHC II^-/-^ mice. Like PEG cells, the numbers of memory OT II cells rapidly declined over time and were undetectable by 150 days post transfer (Fig. S6D-6E). *In vitro* generated memory OT II cells also did not persist in WT mice (Fig. S6G and 6H). Collectively, these observations show that *in vitro* generated memory PEG and OT II cells do not persist upon transfer into wild-type or MHC II^-/-^ mice.

### *In vivo* generated memory PEG CD4^+^ T cells do not persist upon adoptive transfer into MHC II^-/-^ and WT mice

The preceding results show both PEG and OT II memory CD4^+^ T cells generated *in vitro* do not persist in MHC II^-/-^ and WT mice. To determine whether this was due to the *in vitro* generation process, we generated memory PEG cells *in vivo,* transferred them into congenic recipients and monitored their longevity over time (Fig. 5A). Akin to *in vitro* generated memory cells, memory PEG cells generated *in vivo* proliferated and produced IFN-γ in the spleen, dLN and non-dLN of recipient mice (Fig. 5B), and expressed markers characteristics of Tem and Tcm cells (Fig. 5C). Similarly, *in vivo* generated memory PEG cells generated also declined in both MHC II^-/-^ (Fig. S7A, Fig. 5D and 5E) and WT (Fig. S7B, Fig. 5F and 5G) mice such that they were undetectable by 120 days post-transfer.

**Fig. 5.**
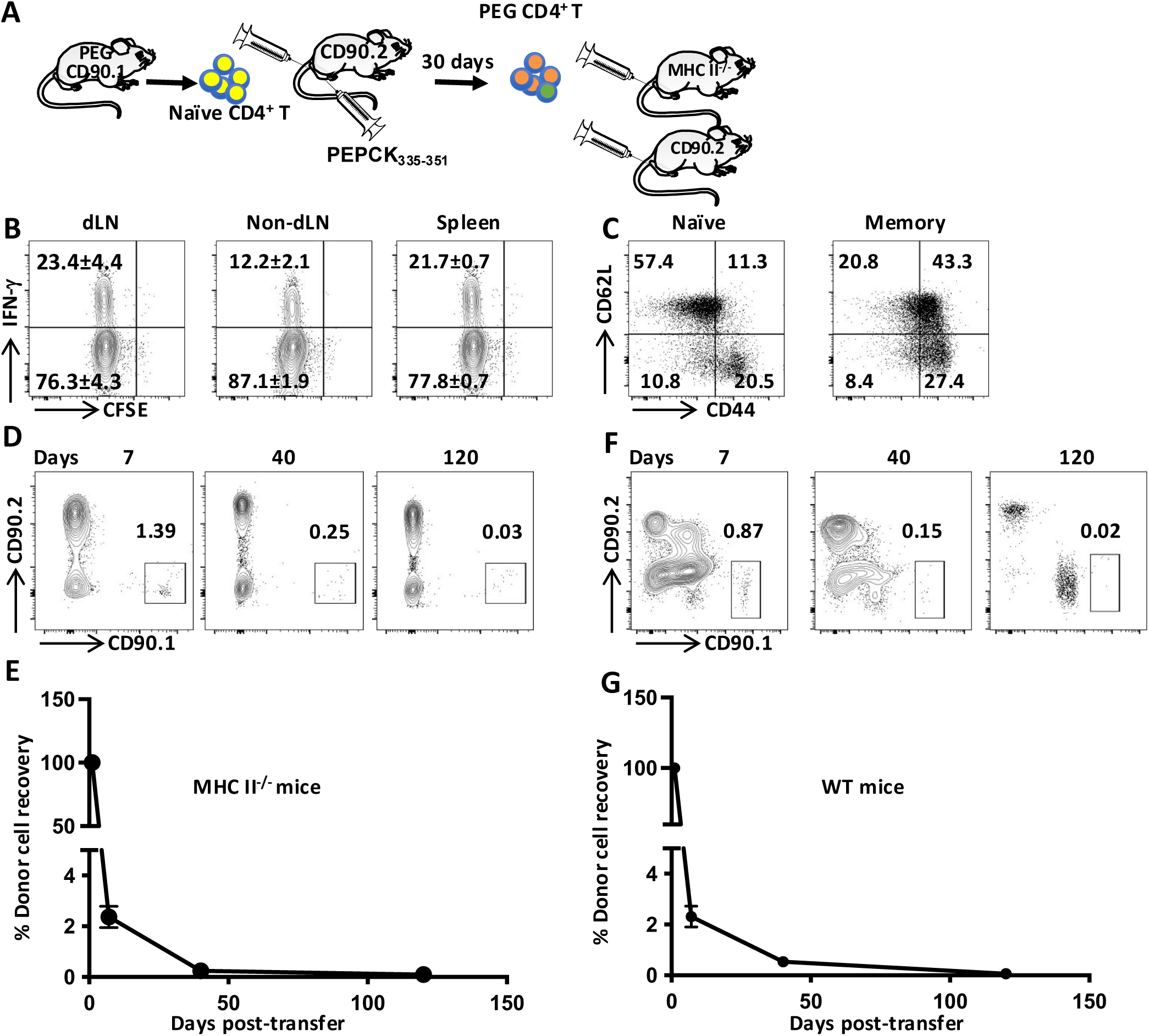
Memory PEG cells generated *in vivo* do not persist following adoptive transfer into MHC II^-/-^ and WT mice. (**A**) Schematic protocol for generating memory PEG cells *in vivo*. Naïve CD4^+^ PEG cells (1 x 10^6^) were transferred intravenously into congenic mice that were injected with 5 nmol PEPCK_335-351_ and 25 μg CpG the next day. Cell proliferation and IFN-γ production by PEG cells in dLN, non-dLN and spleen of recipient mice were assessed in 7 days (**B**). After 30 days, mice were sacrificed and the donor transgenic cells were isolated and assessed for expression of CD62L and CD44 molecules. (**C**) One million (10^6^) purified memory PEG cells were injected intravenously into MHC II^-/-^ (**D** and **E**) or WT mice (**F** and **G**) and their numbers were determined at the indicated days by tetramer enrichment technique.

### Memory PEG CD4^+^ T cells activated *in vivo* with PEPCK_335-351_ peptide do not persist in the absence of PEPCK antigen

Memory PEG CD4^+^ T cells generated *in vitro* or *in vivo* do not persist upon adoptive transfer into MHC II^-/-^ and WT mice. To determine if the adoptive transfer process impacted their survival, we monitored the longevity of PEG cells activated *in vivo* in recipient mice (Fig. 6A). We transferred naïve PEG cells into congenic mice, activated them by injecting PEPCK_335-351_ the next day and assessed their longevity over time.by tetramer enrichment technique ^21^. Transferred PEG cells expanded about 2-3 folds within 1 week and then contracted in about 4 weeks (90% reduction compared to 1 week) (Fig. 6B, Fig. S8A and 8B), and were completely undetectable by 280 days post-transfer (Fig. 6B and Fig. S8B). Similarly, endogenous (recipient) PEPCK-specific CD4^+^ T cells also expanded and contracted akin to PEG cells following injection of PEPCK peptide and their numbers declined over time (Fig. 6C and Fig. S8C).

**Fig. 6.**
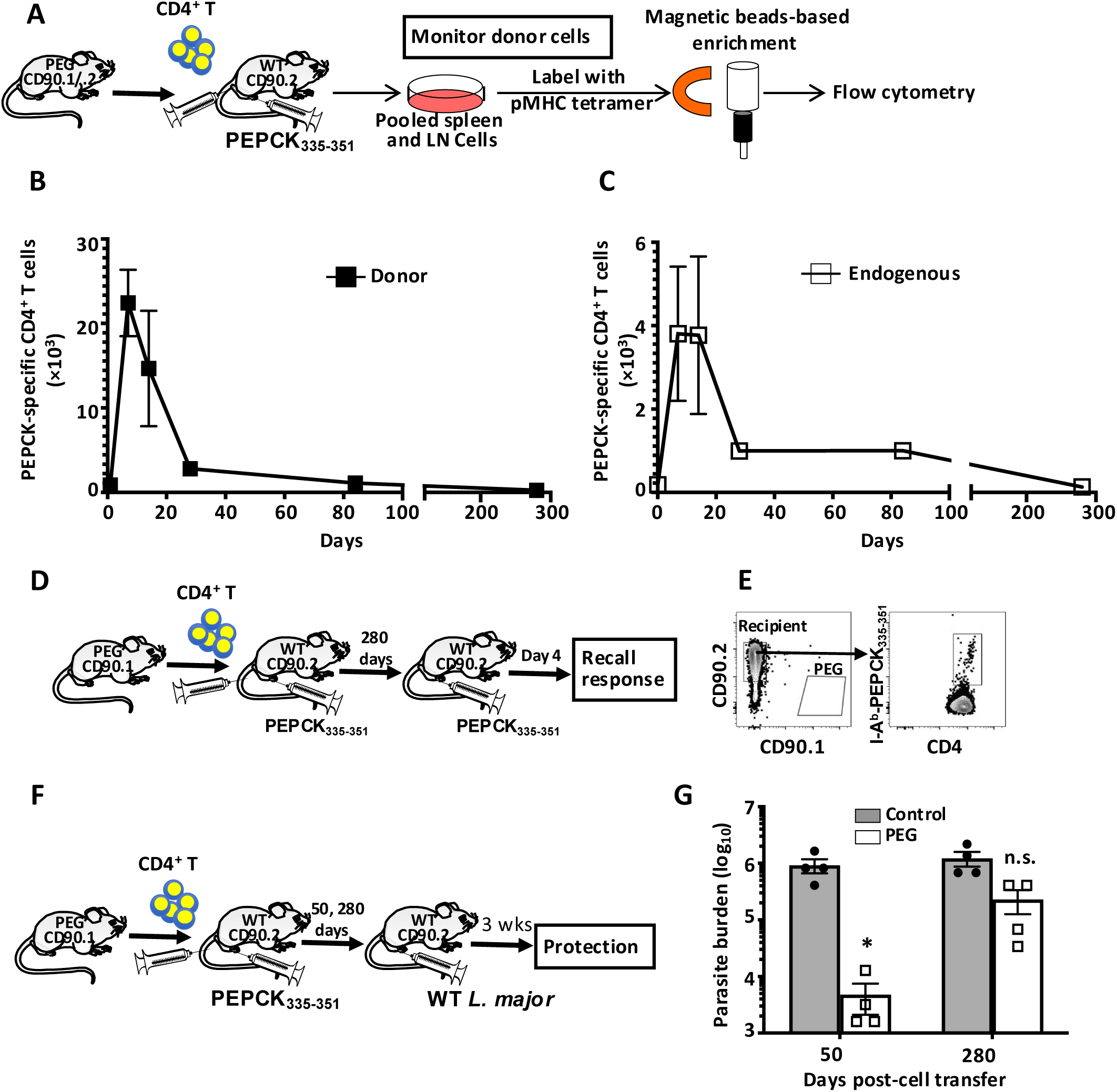
Memory PEG CD4^+^ T cells generated *in vivo* (without adoptive transfer) do not persist in absence of PEPCK antigen. (**A**) Schematic protocol for monitoring memory PEG cells without adoptive transfer in the mice. Ten thousand (10^4^) naïve PEG cells were adoptively transferred into congenic C57BL/6 mice that were then injected with 5 nmol PEPCK_335-351_ and 25 μg CpG in the footpad the next day. At the indicated times, donor PEG cells (**B**) and endogenous (recipient) PEPCK-specific CD4^+^ T cells (**C**) in the spleens and all peripheral lymph nodes were assessed by magnetic bead-based tetramer enrichment. (**D**) Schematic protocol for assessing recall responses in recipient mice. Some recipient mice were rechallenged at 280 days with 5 nmol PEPCK_335-351_ and 25 μg CpG and after 4 days, donor PEG (CD90.1^+^) and endogenous (CD90.2^+^) CD4^+^ T cells were assessed by magnetic beads-based enrichment using I-A^b^-PEPCK_335-351_ tetramer (**E**). (**F**) Schematic protocol for assessing protection in recipient mice. Some recipient mice and age-matched naïve controls were challenged with 10^6^ virulent *L. major* and parasite burden was assessed after 3 weeks post-challenge. Results presented are representative of 2 independent experiments (n = 4 mice in each group per experiment) with similar results. *, p < 0.05; n.s., not significant.

To ensure that lack of detection of PEG cells was not due to poor sensitivity of the enrichment method, we challenged the recipient mice with PEPCK_335-351_ in CpG in a recall response (Fig. 6D). Such recall challenge did not lead to detection of PEG CD4^+^ T cells in the recipient mice (Fig. 6E). In contrast, endogenous (recipient) PEPCK-specific CD4^+^ T cells were detectable in the same mice (Fig. 6E). We believe that this is most likely due to expansion of newly produced naïve endogenous PEPCK-specific CD4^+^ T cells. Furthermore, this tetramer enrichment method can detect as few as 100 PEG cells transferred into congenic recipient mice the next day or within 1 week following PEPCK peptide challenge (Fig. S8D), validating its high sensitivity. The disappearance of memory PEG cells was associated with loss of protection against virulent *L. major* challenge (Fig. 6F and 6G). Collectively, these findings show that the life span of *Leishmania*-specific memory CD4^+^ T cells in the absence of its cognate antigen.

### Persistent of *Leishmania* major parasites is critical for maintenance of memory PEG CD4^+^ T cells

The preceding data showed that *Leishmania*-specific memory CD4^+^ T cells generated either *in vitro* or *in vivo* do not persist when transferred into an antigen deficient host, suggesting that *Leishmania* antigen may be required for their persistence. To test this, we adoptively transferred PEG cells into congenic mice and challenged them with WT *L. major*, which persists indefinitely or *dhfr-ts^-^ L. major* (Fig. 7A), which induces comparable early protection like WT parasites (Fig. 1A-C) but are completely cleared by 3 months post infection ^16^. As expected, both *dhfr-ts^-^* and WT *L. major* induced similar degree of proliferation and IFN-γ response in PEG cells early after challenge (Fig.7B, Fig. S9A and 9B). However, by 5 weeks post challenge, PEG cells continued to expand in the WT *L. major* challenged group but declined in those challenged with *dhfr-ts^-^ L. major* (Fig. 7B and Fig. S9C). Thereafter, PEG cells declined in recipient mice infected with *dhfr-ts^-^ L. major* and became undetectable by 100 days. In contrast, although the numbers of PEG cells also declined in mice infected with WT *L. major*-infected, the remained detectable up to 250 days post-infection (Fig. 7B and Fig. S9C).

**Fig. 7.**
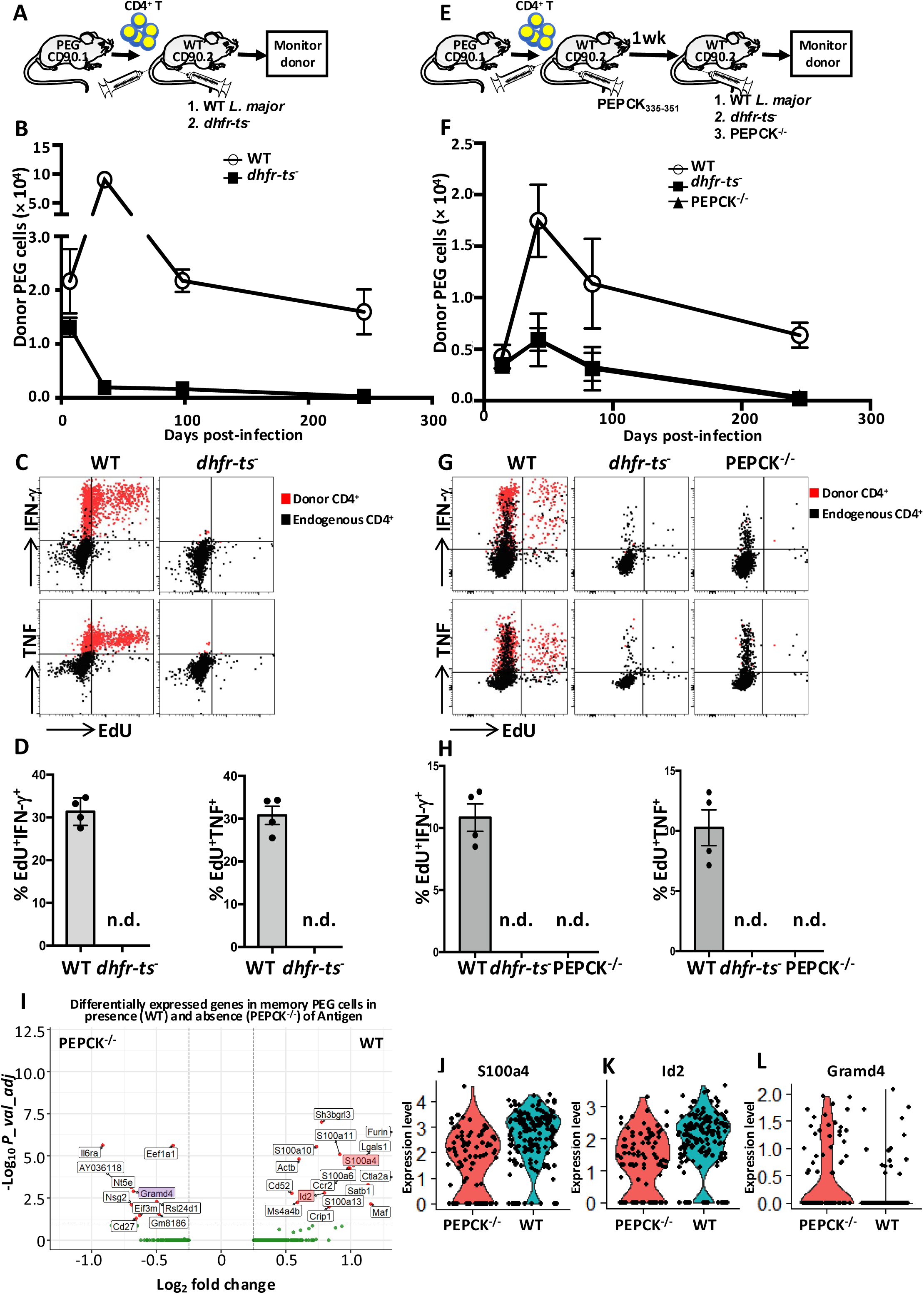
Antigen-specific memory PEG CD4^+^ T cells persist only in the presence of their cognate antigen. (**A**) Schematic protocol for data presented in Figures B-D. Ten thousand (10^4^) naïve PEG cells were transferred into congenic mice that were then infected with either 10^6^ WT or *dhfr-ts^-^ L. major* in the footpads the next day. At the indicated times, donor PEG cells numbers were determined by flow cytometry (**B**). (**C** and **D**) Recall responses in recipient mice infected with WT or *dhfr-ts^-^ L. major*. At 280 days after transfer and challenge with parasites, mice were rechallenged with WT *L. major*, sacrificed after 4 days and donor PEG cells were enriched by magnetic beads enrichment technique. The enriched cells were stimulated overnight with splenic DCs pulsed with PEPCK_335-351_ peptide in the presence of EdU and proliferation (EdU^+^) and production of cytokines (IFN-γ and TNF) by donor PEG and endogenous cells were assessed (**C**) and represented graphically as percentage of EdU^+^IFN-γ^+^ or EdU^+^TNF^+^ cells (**D**). (**E**) Schematic protocol for data presented in Figures F-H. Ten thousand (10^4^) naïve PEG cells were transferred into congenic mice that were then injected with 5 nmol PEPCK_335-351_ and 25 μg CpG in the footpads the next day. After 1 week, mice were then infected with either 10^4^ WT, *dhfr-ts^-^* or PEPCK^-/-^ *L. major* and at indicated times, donor PEG cells were monitored by flow cytometry (**F**). (**G** and **H**) recall responses were measured in WT, *dhfr-ts^-^* or PEPCK^-/-^ *L. major-*infected mice using the same method as above. The production of cytokines (IFN-γ and TNF) and proliferation by donor PEG cells and endogenous CD4^+^ cells were assessed (**G**) and represented graphically as percentage of EdU^+^IFN-γ^+^ or EdU^+^TNF^+^ cells (**H**). (**I**-**L**) Differentially expressed genes in memory PEG cells isolated from mice that express or do not express the cognate antigen (persistent WT or PEPCK^-/-^ *L. major*, respectively). PEG cells were transferred into congenic mice, injected with PEPCK_335-351_ plus CpG and infected with either WT or PEPCK^-/-^ *L. major* after 1 weeks. After 50 days, mice were sacrificed, and PEG cells were isolated and used for scRNAseq. (**I**) Volcano plots depicting fold changes of differentially expressed genes in memory PEG CD4^+^ T cells in the presence (WT *L. major*) or absence (PEPCK^-/-^ *L. major*) of their cognate antigen. *p < 0.05 for all genes. Violin plots of S100a4 (**J**), Id2 (**K**) and Gramd4 (**L**) expression in memory PEG CD4^+^ T cells in the presence (WT *L. major*) or absence (PEPCK^-/-^ *L. major*) of antigen. Results presented are representative of 2 independent experiments (n = 4 mice per group) with similar results.

To confirm loss of PEG cells in the group infected with non-persistent *dhfr-ts^-^ L. major*, we rechallenged both groups with WT *L. major* and assessed cell expansion and cytokine response after 4 days (Fig. 7C). Donor PEG cells in mice infected with WT *L. major* expanded (proliferated) and produced cytokines (IFN-γ and TNF) in response to *L. major* challenge (Fig. 7C and D). In contrast, PEG cells remained undetectable in the mice infected with *dhfr-ts^-^ L. major* (Fig. 7C and D), suggesting that memory PEG CD4^+^ T cells were lost in the absence of *Leishmania* antigen.

### Expression of cognate (PEPCK) antigen by persisting parasites is critical for maintenance of memory PEG CD4^+^ T cells *in vivo*

It is conceivable that persistence of memory PEG CD4^+^ T cells in mice infected with WT (but not *dhfr-ts^-^*) *L. major* may be due to resulting inflammatory environment created by persisting parasites and unrelated to antigen (PEPCK) persistence. To test this, we used PEPCK^-/-^ *L. major* which persist indefinitely in infected mice similar to WT *L. major* ^22^ but lack PEPCK (antigen) expression. We transferred PEG cells into naïve congenic mice, challenged them with PEPCK peptide the next day and after 1 week, infected them with WT, *dhfr-ts^-^* or PEPCK^-/-^ *L. major*. We then compared the longevity of memory PEG CD4^+^ T cells in these mice by the tetramer enrichment technique (Fig. 7E). At 2 weeks post-infection, the numbers of PEG cells were similar in all groups (Fig. 7F and Fig. S9D and 9E), confirming that PEG cells were activated to the same magnitude in all groups following PEPCK peptide injection. Thereafter, PEG cells expanded by more than 3 folds by 6 weeks post-infection in mice infected with WT *L. major* and their absolute numbers remained relatively constant up to 250 days. In contrast, the number of PEG cells in mice infected with *dhfr-ts^-^* (which do not persist) or PEPCK^-/-^ (which persist but does not express PEPCK antigen) parasites declined rapidly and were undetectable by 250 days post-transfer (Fig. 7F and Fig. S9E). As expected, both WT and PEPCK^-/-^ (but not *dhfr-ts^-^*) parasites were detected at the infection sites in similar numbers by 250 days post infection (Fig. S9F).

To confirm the loss of PEG cells in mice injected with *dhfr-ts^-^* or PEPCK^-/-^ parasites, we rechallenged them with WT *L. major* and assessed *in vivo* recall response after 4 days by flow cytometry. We observed proliferation and cytokine (IFN-γ and TNF) production by PEG cells only in the mice infected with WT, but not *dhfr-ts^-^* or PEPCK^-/-^ *L. major* (Fig. 7G and 7H). Collectively, these findings confirm that mere persistence of parasites and the resulting inflammatory environment were not sufficient to maintain memory CD4^+^ T cells *in vivo*. They show that the presence of cognate antigen is obligatory for long-term maintenance of *Leishmania*-specific memory CD4^+^ T cells *in vivo*.

To understand the possible molecular mechanisms why cognate antigen is critical for maintenance of *Leishmania*-specific CD4^+^ memory T cells, we conducted scRNA-seq analysis on purified memory PEG cells isolated from mice infected with WT (persistent antigen) or PEPCK^-/-^ (no antigen) *L. major* (Fig. S10A). We identified 27 differentially expressed genes (Fig. 7I and Fig. S10) of which 17 were upregulated in the presence of antigen (WT *L. major*) while 10 genes were upregulated in absence of antigen (PEPCK^-/-^ *L. major*). Interestingly, S100A4, which has been shown to be involved in TCR signalling ^23,24^, was upregulated in memory PEG cells from WT *L. major*-infected mice (Fig. 7I and Fig. 7J). Also, Id2, whose deficiency in CD4^+^ or CD8^+^ T cells leads to increased cell death or apoptosis ^25,26^, was upregulated in memory PEG cells from WT *L. major*-infected mice (Fig. 7I and Fig. 7K). In contrast, GRAMD4, also known as death-inducing protein mediating apoptotic function ^27^, was upregulated in memory PEG cells from PEPCK^-/-^ *L. major*-infected mice (Fig. 7I and Fig. 7L).

### Endogenously generated PEPCK-specific memory CD4^+^ T cells do not persist in the absence of PEPCK antigen

The preceding observations focused on assessing longevity of *in vitro* and *in vivo* generated memory transgenic (PEG) CD4^+^ T cell in the presence or absence of its cognate antigen. Because transgenic cells are monoclonal, we investigated long-term survival of endogenously PEPCK-specific memory CD4^+^ T cells that polyclonal in nature with different TCR affinities. Following PEPCK peptide challenge (Fig. 8A), or infection with *dhfr-ts^-^ L. major* (data not shown), endogenous PEPCK-specific CD4^+^ T cells massively expanded in numbers (Fig. S11, Fig. 8B and 8C) and this was associated with corresponding increase in mean fluorescence of intensity (MFI) of CD44 expression (Fig. 8D). As expected, the expansion was followed by contraction and decline such that by 350 days post-challenge, the numbers of PEPCK-specific cells were similar to those in naïve mice (Fig. 8C). Interestingly, the MFI of CD44 expression on PEPCK-specific CD4^+^ T cells at 350 days was like their naïve counterparts (Fig. 8D), suggesting that the PEPCK-specific CD4^+^ T cells detected at 350 days were naïve cells that could have arisen from normal hematopoiesis. To confirm this, we performed secondary peptide challenge at 50 or 350 days and compared expansion of PEPCK-specific CD4^+^ T cells in spleens and LNs after 3 days. PEPCK-specific CD4^+^ T cells massively expanded and produced effectors cytokine (IFN-γ) following peptide rechallenge at 50 days (when PEPCK antigen was still present) but not at 350 days (Fig. 8E-H). These observations confirm that the CD44^lo^ PEPCK-specific cells we detected in these mice at 350 days were indeed newly generated naïve cells. Collectively, these findings indicate that the maintenance of *Leishmania*-specific CD4^+^ memory T cells (both transgenic and endogenous) is dependent on the continuous presence of *Leishmania* antigen.

**Fig. 8.**
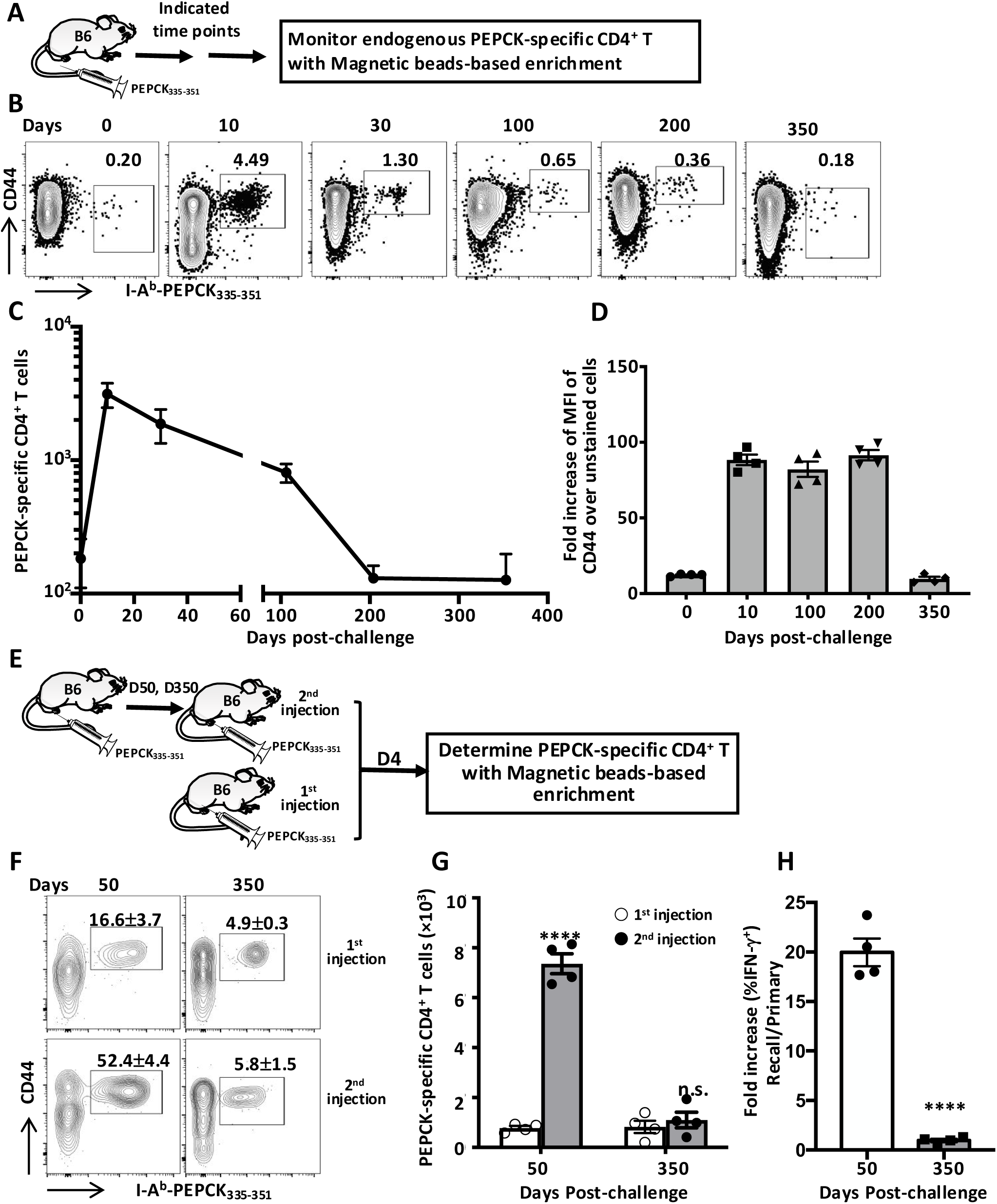
Loss of endogenous PEPCK-specific memory CD4^+^ T cells in the absence of antigen. (**A**) Schematic protocol to assess longevity of endogenously generated PEPCK-specific memory CD4^+^ T cells. C57BL/6 mice were injected with 5 nmol PEPCK_335-351_ and 25 μg CpG and sacrificed at the indicated times to assess the numbers of PEPCK-specific CD4^+^ T cells by magnetic bead tetramer enrichment technique. The percentage (**B**) and absolute numbers (**C**) of activated (CD44^+^) PEPCK-specific T cells were calculated (**C**). **(D)** Fold increase in the mean fluorescence intensity (MFI) of CD44 expression on PEPCK-specific CD4^+^ T cells at different times after peptide challenge over unstained controls). **(E)** Schematic protocol for measuring recall responses to confirm loss of endogenous PEPCK-specific CD4^+^ memory T cells. At the indicated times (50 and 350 days) after antigen injection, mice (together with some age-matched naïve controls, 1^st^ PEPCK injection) were rechallenged with PEPCK peptide plus CpG and the percentages (**F**) and absolute numbers (**G**) of PEPCK-specific CD4^+^ T cells were determined after magnetic bead-based tetramer enrichment. In addition, the fold increase in the percentage of IFN-γ^+^ cells over naïve controls was assessed by flow cytometry (**H**). Results presented are representative of 2 independent experiments (n = 4 mice per group per experiment) with similar results. ****, p < 0.0001; n.s., not significant.

## DISCUSSION

Antigen recognition via TCR is required for activation of naïve T cells that leads to the generation of effective T cell-mediated immunity against infectious agents. In addition, continuous antigen recognition is thought to be required throughout the different stages of CD4^+^ T cell response including expansion, differentiation, acquisition of effector functions and generation of memory ^28^. Indeed, extended antigen presentation by DCs is important for sustaining differentiation of CD4^+^ T cells into memory cells ^29^. This is different from CD8^+^ T cells which require only a short period of antigen presentation for priming, activation and differentiation into effector and memory cells ^30,31^. Thus, continuous antigen presentation and antigen persistence are essential for eliciting optimal effector CD4^+^ T cell responses ^32^. Here, we show that extended antigen presence is also crucial for maintenance of memory CD4^+^ T cells *in vivo*.

A strong CD4^+^ T cell response is critical for providing both primary and secondary (memory) protection against cutaneous leishmaniasis in both human and mice. The mechanisms and pathways leading to the development of effector and memory CD4^+^ T cells in *Leishmania*-infected mice have been studied and reviewed extensively ^11,12,33–35^. These studies show that recovery from cutaneous leishmaniasis in both human and mice is associated with resistance to subsequent infections, known as infection-induced immunity ^11,12,35^. This infection-induced immunity, which is mediated by *Leishmania*-specific CD4^+^ T cells ^12,17,18,35^, is believed to be dependent upon parasite persistence at the primary infection sites because infection protocols or manipulations that lead to complete parasite clearance results in loss of immunity ^14,15^. In line with this, Leishmanization, which is the inoculation of few live *Leishmania* in a hidden part of the body ^36^, and vaccination with genetically attenuated but persistent parasites ^37^ have been shown to provide long term protection against *Leishmania* infections. In contrast, vaccination with killed parasites ^38^, subunit antigens ^39,40^ or non-persistent genetically modified parasites ^16^ do not provide long term protection. These findings strongly indicate that maintenance of *Leishmania*-specific memory CD4^+^ T cells that mediate secondary (memory) immunity may be dependent on the presence of residual parasites (antigens). In line with this, we found that mice infected with *dhfr-ts^-^ L. major* were protected against virulent *L. major* challenge when *dhfr-ts^-^* parasites were still detectable in these mice, but this protection was lost following complete clearance of parasites. This loss of protection was associated with loss of polyclonal and antigen (PEPCK)-specific CD4^+^ T cells that are associated with secondary protection in the challenged mice.

Although the prevailing opinion is that maintenance of memory CD8^+^ T cells is antigen independent ^4^, whether memory CD4^+^ T cells are maintained in the absence of antigen is controversial. Hu et al showed that CD4^+^ effector T cells generated *in vitro* transitioned into memory cells upon adoptive transfer into MHC class II^-/-^ mice and these memory cells remained stable for about 50-90 days after transfer ^6^. In contrast, other studies showed that not all CD4^+^ memory T cells are long lived ^41^. For example, CD4^+^ memory T cells have been shown to gradually decline in mice infected with LCMV ^4^, Listeria ^5^, Salmonella ^42^ and other antigens ^43,44^.

Most of the studies that investigated longevity of antigen-specific memory CD4^+^ T cell responses used model antigens, such as OVA, 2W peptide and others ^43,45,46^. These model antigens may not reflect natural features of antigens from pathogens that co-evolved with their hosts. To overcome this limitation, we exploited the immunodominant properties of *Leishmania* antigen, PEPCK ^18^. Our identification of an immunodominant epitope of PEPCK led to the development of a tetramer to enrich PEPCK-specific CD4^+^ T cells, cloning of their TCR genes and generation of first *Leishmania*-specific CD4^+^ TCR transgenic (PEG) mice on a C57BL/6 background, which is the murine model of cutaneous leishmaniasis that closely mimic human disease. These PEG mice provided a unique and meaningful tool to study *Leishmania*-specific CD4^+^ T cell responses in a manner never done previously.

To assess longevity of *Leishmania*-specific memory CD4^+^ T cells in the absence of antigen, we first generated PEG memory CD4^+^ T cells *in vitro* ^6^ and monitored their persistence in MHC II^-/-^ or WT mice as was done previously ^6^. Surprisingly and in contrast to previous reports ^6^, the transferred cells were not maintained but rather declined over time as in other publications ^4,5,42^. Interestingly, surviving PEG cells were still functional (produced effector cytokines upon *in vitro* restimulation), which is in line with a previous study ^4^. The decline of memory PEG cells was not due to i*n vitro* generation process because we found that memory PEG cells generated *in vivo* also declined in numbers upon transfer into MHC II^-/-^ or WT mice.

The maintenance of CD4^+^ T cell memory that mediate concomitant immunity during some chronic infectious diseases such as malaria ^47^ and leishmaniasis ^15,17^ is thought to be dependent on the persistence of antigens. Infection with *L. major* induces a strong Th1 response that robustly controls secondary infections at distal sites, while parasites at primary site of infection are not fully cleared ^11,12,34^. When persisting parasites at the primary infection sites are cleared (as is seen in IL-10^-/-^ mice), immunity to reinfection is lost ^14,15^. Recent reports show that antigen-specific CD4^+^ memory T cells decline over time, persisting at a low frequency (about 2 folds more than naïve antigen specific CD4^+^ T cells) following clearance of pathogen or antigen in mice ^4,5,42–44^. Our data do not support these findings, because *Leishmania*-specific TCR transgenic memory CD4^+^ T cells were completely lost in absence of their cognate antigen. Similarly, endogenous *Leishmania* PEPCK-specific memory CD4^+^ T cells generated following infection with *Leishmania* or immunization with PEPCK peptide did not persist in absence of PEPCK antigen.

TCR transgenic mouse is a great tool for assessing antigen-specific T cell responses although it is believed that adoptive transfer of abnormally high numbers of antigen-specific T cells may not correctly represent what happens during infections. This is because such a high number of cells may hinder maximum proliferation and differentiation of these cells upon antigen encounter, thereby negatively impacting on memory T cell generation and function ^48,49^. In the present study, we adoptively transferred between 10^3^ to 10^4^ PEG cells, which is similar to the frequency of natural PEPCK-specific T cells in uninfected naïve mice ^18^. Following transfer and PEPCK peptide challenge, the dynamics of PEG cell activation, expansion and contraction was comparable to endogenous PEPCK-specific CD4 T cells (Fig. 6), suggesting that 10^3^-10^4^ PEG cells responded to their cognate antigen akin to endogenous PEPCK CD4^+^ T cells.

Effective vaccine development and vaccination strategy against any infectious agent require in-depth understanding of the induction and longevity (maintenance) of protective memory T cells against that pathogen. Unfortunately, this is rarely done due to general acceptance of the dogma that maintenance of memory T cells is antigen-independent ^50^. Our study clearly provides evidence that continuous presence of antigen is required for sustaining *Leishmania*-specific memory CD4^+^ T cells. Although there are reports of antigen-independent long-lived vaccinia virus-specific CD4^+^ memory T cells in humans ^51–53^, it is not known whether these cells are maintained in absence of cross-reactive environmental antigens ^41^ or by contact with persisting antigen bound as immune complexes on follicular dendritic cells ^54^. A recent study showed that intercellular transfer of telomeres from APCs rescues T cells from senescence and increases their longevity and ability to confer long-term immune protection ^55^. It would be interesting to determine whether APCs containing peptides derived from persisting antigen could also transfer telomers to their respective antigen-specific CD4^+^ T cells thereby promoting their longevity.

To understand why long-term survival of *Leishmania*-specific memory CD4^+^ T cells require persistence of their cognate antigen, we compared the transcriptional profile of purified memory PEG cells recovered from mice infected with WT or PEPCK^-/-^ *L. major*. Twenty-seven genes were differentially and significantly expressed by the 2 groups of cells, indicating that environmental difference (i.e. presence or absence of cognate antigen) did influence the transcriptome of memory CD4^+^ T cells. Interestingly, *S100A4*, which is one of the highly upregulated genes in PEG cells maintained in the presence of antigen (WT *L. major*-infected group), has been shown to be expressed exclusively by memory CD4^+^ or CD8^+^ T cells ^24^. S100A4 directly binds to Lck and Fyn and reciprocally regulate their kinase activity towards the CD5 ^23^, which is a negative modulator of TCR signalling ^56,57^. Although it is unclear how S100A4 may contribute to long term persistence of memory CD4^+^ T cells, we speculate that this may be related to its effect on TCR signaling. Another gene that was significantly upregulated in in PEG cells maintained in the presence of antigen is Inhibitor of DNA binding 2 (Id2), a helix–loop–helix transcriptional regulator which has been shown control the numbers of effector and memory CD8^+^ T cells^26,58^. Id2-deficient CD4 T cells express higher levels of Bim and SOCS3, resulting to increased cell death in an EAE mouse model^25^. Thus, it is conceivable that the induction of Id2 promotes survival of memory CD4^+^ T cells. In contrast the gene that encodes GRAMD4, also known as death-inducing protein, was upregulated in memory PEG cells maintained in the absence of antigen (i.e. isolated from PEPCK^-/-^ *L. major*-infected mice). GRAMD4 interacts with Bcl-2, promotes Bax mitochondrial relocalization and oligomerization, and is capable of severely disrupting mitochondrial membrane integrity resulting the initiation of cellular apoptosis^27^. Taken together, these observations suggest that antigen-mediated TCR signalling, and cell survival pathways may be involved for longevity of memory CD4^+^ T cells.

Because of long-term nature of the studies reported here, we used female animals exclusively and this precluded the use of male animals because of infighting when male animals are housed together in the same cages. This exclusive use of female animals prevents us from making broad conclusions on longevity of *Leishmania*-specific memory T cells across both sexes and is a limitation of this study. Comparative studies in both male and female recipients will permit investigations into whether biological sex is an important regulator of longevity of *Leishmania*-specific memory CD4^+^ T cells and is our current investigative focus.

## MATERIALS AND METHODS

### Mice

Six to 8 weeks old female C57BL/6 (B6), Ly5.1(CD45.1^+^) and OT II mice were obtained from Charles River, St Constante PQ, Canada. B6 Thy1.1 and B6 MHC II^-/-^ mice were purchased from The Jackson Laboratory. PEPCK-specific TCR Tg PEG mouse founders were originally generated on C57BL/6 background. The PEG founders were bred with B6 mice for the first 5 generations and then PEG males were bred with B6 Thy1.1 WT mice at the University of Manitoba Central Animal Care Facilities. All mice were housed at the University of Manitoba Central Animal Care Facilities and were maintained according to the recommendations of the Canadian Council of Animal Care.

### Infection protocol and parasite quantification

*Leishmania major* wildtype (WT), *dhfr-ts* deficient (*dhfr-ts^-^*), and PEPCK deficient (PEPCK^-/-^) strains (on MHOM/80/Fredlin strain) were grown in M199 culture medium (Sigma, St. Louis, MO) supplemented with 20% heat inactivated FBS, 2 mM glutamine, 100 U/ml penicillin, and 100 μg/ml streptomycin (thymidine at 10 μg/ml was added for *dhfr-ts^-^* cultures). For infection, mice were injected with one million (1 x 10^6^) 7-day stationary-phase promastigotes in 50 μl PBS suspension into the left hind footpad. Lesion sizes were monitored by measuring footpad swelling with calipers. For intradermal infection, mice were injected with 10^6^ promastigotes in 10 μl PBS suspension into the ear. For measuring secondary (recall) protection, mice were challenged with 10^6^ virulent *L. major* promastigotes in the contralateral footpads. Lesion size was monitored weekly and parasite burden was measured by limited dilution ^18,59,60^ at 3 weeks after euthanasia.

### Assessment of infection-induced immunity

Six to 8 weeks old female C57BL/6 were injected in the right hind footpads with either PBS, 10^6^ WT *L. major*, *dhfr-ts^-^* or *dhfr-ts*^-^ mixed with 10^4^ WT *L. major*. After 6 and 24 weeks, the mice (together with their age-matched controls) were challenged in the left hind footpads with 10^6^ WT *L. major*. Three days after challenge, footpad swelling (delayed-type hypersensitivity, DTH response) in the challenged footpads was measured with digital calipers. The challenged mice were sacrificed after 3 weeks to determine parasite burden.

### Assessment of longevity of CD4^+^ T cells that mediate infection-induced immunity by adoptive transfer

Six to 8 weeks old female CD90.1 C57BL/6 were injected in the right hind footpads with PBS (naïve) or 10^6^ WT *L. major* and allowed to heal (>12 weeks post-infection). CD4^+^ T cells were purified from naïve controls or healed mice and adoptively transferred (5 x 10^6^ cells) into congenic (CD90.2) recipient mice. At different times, recipient mice were sacrificed to assess the frequency of PEPCK-specific CD4^+^ T cells within the donor cell populations. Some recipient mice were challenged with 10^6^ *L. major* and DTH response was assessed after 3 days. Three weeks after challenge, mice were sacrificed to determine parasite burden by limiting dilution ^18,59,60^.

### Generation of PEG mice

To amplify PEPCK-specific TCR beta and alpha genes, C5BL/6 mice were infected with 5 x 10^6^ *L. major* in the hind footpads. After 5 weeks, infected mice were sacrificed and splenocytes were stained with I-A^b^-PEPCK_335–351_ tetramer. PEPCK-specific TCR beta and alpha genes were amplified from tetramer binding cells by single-cell RT-PCR. The retroviral vector pMXs-TCRβ-P2A-TCRα-IRES-GFP was used to express TCRα and TCRβ proteins and to validate function and tetramer binding of the TCR. The TCR alpha and beta genes used for generating PEG mice (TCR31) were identified as TRBV13-2-J2-3-D2 and TRAV6-J17 with the IMGT/V-Quest tool (http://www.imgt.org/). Then complete TCRβ-P2A-TCRα was subcloned into VA hCD2 transgenic cassette vectors ^19^, which uses the human CD2 promoter to drive the expression of PEPCK-specific TCR in T cells. Linearized transgenic plasmid DNA fragments containing the TCR cDNAs with Sca I were injected into the pronucleus of fertilized C57BL/6 eggs. And then injected eggs were implanted into surrogate mothers to obtain offspring. Founders and their progeny were screened by genotyping and confirmed by flow cytometry using I-A^b^-PEPCK_335–351_ tetramer staining. TCR Tg founder number 27 was selected and used in all experiments presented in this paper.

### Characterization of PEG mice

For *in vitro* co-culture, purified and CFSE-labelled naïve PEG CD4^+^ T cells (2 x 10^5^ cells/ well) were stimulated with splenic DCs (DC:T ratio 1:10) in presence of PEPCK_335-351_ peptide (0.2, 2, 20 μM) or 50 μg/ml SLA in 96-well round-bottom plates for 4 days. For *in vivo* studies, CFSE-labelled PEG or equal numbers PEG and polyclonal CD4^+^ T cells (5 x 10^5^) were adoptively transferred into congenic C57BL/6 mice. On next day, the recipient mice were challenged with 5 x 10^6^ WT or PEPCK^-/-^ *L. major* in the hind footpads. After 4 days, mice were sacrificed and cell proliferation and IFN-γ production by the donor PEG cells were assessed by flow cytometry.

### MP-IVM analysis of DC:T cell dynamics in the popliteal lymph node

Dendritic cells, naïve polyclonal or PEG TCR transgenic CD4^+^ T cells were labeled with Celltracker Green (CMFDA; 2 μM), Celltracker Blue (CMAC; 15 µM) or Celltracker Orange (CMTMR, 20 µM) for 15 min at 37 °C prior to injection as previously described ^61^ for analysis of DC:T cell dynamics in the popliteal lymph node. Five hundred thousand (5 × 10^5^) uninfected, *L. major*-infected or PEPCK_335-351_-pulsed DCs were injected into the hind footpad of C57BL/6 mice. Twelve hours after DC transfer, mice were injected with purified 3 x 10^6^ labelled naïve WT or PEG CD4^+^ T cells.

For Intravital microscopy, mice were anaesthetized and the popliteal lymph nodes microsurgically exposed as previously described ^62^. Imaging depth was typically 80–200 μm below the lymph node capsule. A multiphoton microscope with two Ti-sapphire lasers (Coherent) was tuned to between 780 and 920 nm for optimized excitation of the fluorescent probes used. For four-dimensional recordings of cell migration, stacks of 12 optical sections (512 x 512 pixels) with 4 µm z-spacing were acquired every 15 seconds to provide imaging volumes of 44 μm in depth. Emitted light was detected through 460/50 nm, 525/70 nm and 595/50 nm dichroic filters with non-descanned detectors. All images were acquired using the 20X 1.0 N.A. Olympus objective lens (XLUMPLFLN; 2.0mm WD). Automated 3D tracking of T cell centroids was performed for cell motility analyses using Imaris 8.0. Further cell track parameters (arrest coefficient and mean displacement) were analyzed in Matlab (Mathworks) as previously described ^63^.

### Enrichment of PEPCK-specific CD4^+^ T cells

Enrichment of antigen-specific (PEPCK-specific) CD4^+^ T cells was performed by a magnetic bead-based procedure as previously described ^21^. Briefly, single cell suspension from all the peripheral lymph nodes and spleens of *L. major*-infected or healed mice were stained with allophycocyanin (APC)-labeled pMHC tetramer to a final concentration of 10 nM for 30 min at 37 °C. After washing with cold cell sorter buffer (2% FBS and 1mM EDTA in PBS), tetramer-labeled cells were resuspended in 200 μl sorter buffer and incubated with 50 μl of anti-APC conjugated magnetic microbeads (Miltenyi Biotec) for 30 min at 4 °C. Thereafter, tetramer-binding cells were enriched by passing the cells through Miltenyi LS column held by a magnet.

### Generation of memory PEG cells in vitro for adoptive transfer

PEG CD4^+^ T cells were stimulated with LPS pre-matured BMDCs (1μg/ml LPS overnight) in the presence of 5 μM PEPCK_335-351_ under Th1 differentiating condition (20 ng/ml rIL-12, 1μg/ml anti-IL-4 antibody and 30 U/ml rIL-2) at 1:100 DC to T cell ratio for 5 days. Thereafter, the DCs were removed by CD11c positive selection kit (StemCell Technology) and the remaining T cells were cultured in fresh complete DMEM medium with 30 U/ml rIL-2 for 5 days. Thereafter, T cells were rested for another 5 days in complete DMEM medium without rIL-2 and assessed for viability, cell surface expression and cytokine production and before cell transfer.

### Generation of memory PEG cells in vivo for adoptive transfer

One million purified PEG CD4^+^ T cells were transferred into congenic C57BL/6 mice and injected with 5 nmol PEPCK_335-351_ mixed with CpG (25 μg) in footpads the next day. After 30 days, PEG cells were enriched from the spleens and all peripheral lymph nodes by cell sorting and assessed for cell surface expression and cytokine production. One million (10^6^) purified memory PEG cells were injected intravenously into MHC II^-/-^ or WT mice.

### In vivo recall responses

For *in vivo* recall response, mice were injected with either 5 nmol PEPCK_335-351_ and 25 μg CpG or 10^6^ *L. major* for 4 days. PEG or endogenous PEPCK-specific CD4^+^ T cells in spleens and all peripheral lymph nodes were enriched using magnetic bead-based method with I-A^b^-PEPCK_335–351_ tetramer. The number of PEPCK-specific CD4^+^ T cells was determined by flow cytometry using counting beads (Biolegend). For measuring cell proliferation or cytokine production, the enriched cells were co-cultured with splenic DCs in the presence of PEPCK_335–351_ peptide and 10 μM EdU overnight. EdU staining was performed according to the manufacturer’s suggested protocol (Invitrogen).

### scRNA-seq analysis

Ten thousand (10^4^) purified PEG CD4^+^ T cells were transferred into congenic mice that were then injected with 5 nmol PEPCK_335-351_ and 25 μg CpG in the footpads the next day. After 1 week, the mice were then infected with either 10^4^ WT or *PEPCK^-/-^* (KO) *L. major* in the footpads. Donor PEG cells were purified at 3 months post-infection by cell sorting. Single-cell cDNA libraries were generated using 10X Genomics Chromium Single Cell 3’ Reagents v3 and Chromium controller. Quality control of the 10× Genomics-indexed cDNA libraries were performed using 2100 Bioanalyzer (Agilent Technologies). The cDNA libraries of PEG cells isolated from WT or PEPCK^-/-^ *L major*-infected mice were pooled together, sequenced on a NovaSeq6000 at Genome Quebec (Montreal, QC, Canada). The Fastq reads were processed with 10× CellRanger software to generate count matrices. Matrices were loaded into the R package Seurat (v3.0.0) for analysis and filtered to remove cells with high mitochondrial content (>5%) and number of RNA molecule (>10000), low RNA counts (<200 features) and ribosome content (<5%), and CD90.1 and CD4 reads were 0. Data were normalized and variable features identified, followed by data scaling and dimensionality reduction by PCA using the identified variable genes. Data were visualized using tSNE.

### Statistical analysis

Data are presented as means ± standard error of mean (SEM). Two-tailed Student’s t-test was used to compare means and SEM between groups using GraphPad Prism software. Differences were considered significant at p < 0.05.

## Supplementary Materials

### Supplementary Figures

**Fig. S1.**
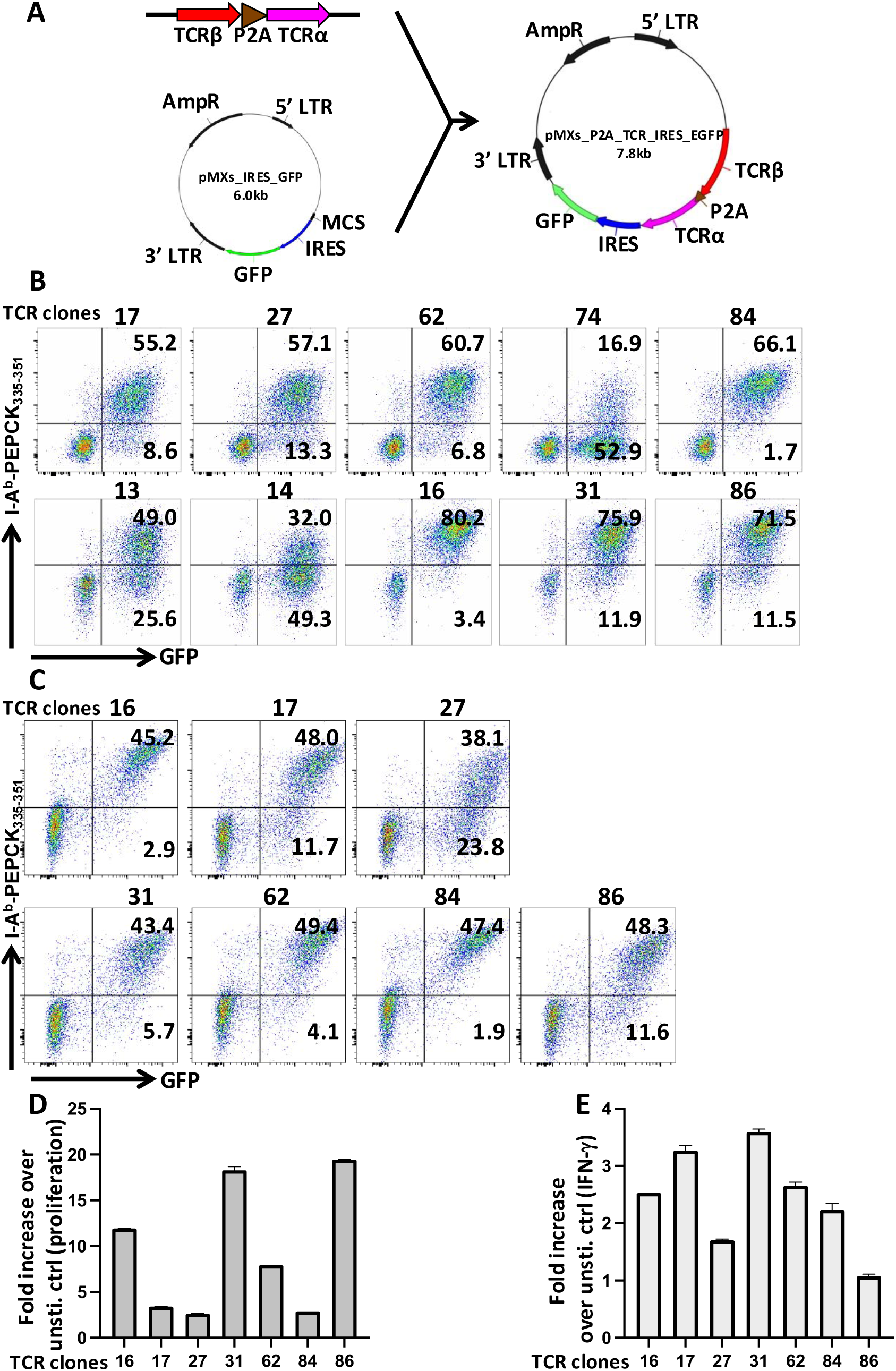
Selection of TCR31 clone for generating TCR Tg mice. (**A**) Schematic protocol for constructing the retroviral vector pMXs-TCRβ-P2A-TCRα-IRES-GFP for expressing TCRα and TCRβ chains. Murine CD4^+^ TG40 cell line (**B**) or primary CD4^+^ T cells (**C**) were infected with recombinant retroviruses expressing PEPCK-specific TCR clones. The antigen specificity of the cloned TCRαβ pairs was analyzed by flow cytometry. The primary CD4^+^ T cells expressing PEPCK-specific TCR were stimulated with splenic DCs pulsed with or without 5 mM PEPCK_335-351_ peptide overnight in the presence of EdU. Cell proliferation and IFN-γ production by GFP+Tet+ CD4+ cells were assessed by flow cytometry. Fold increase in proliferation (**D**) or percentage of IFN-γ (**E**) over unstimulated control (without PEPCK peptide) were analyzed.

**Fig. S2.**
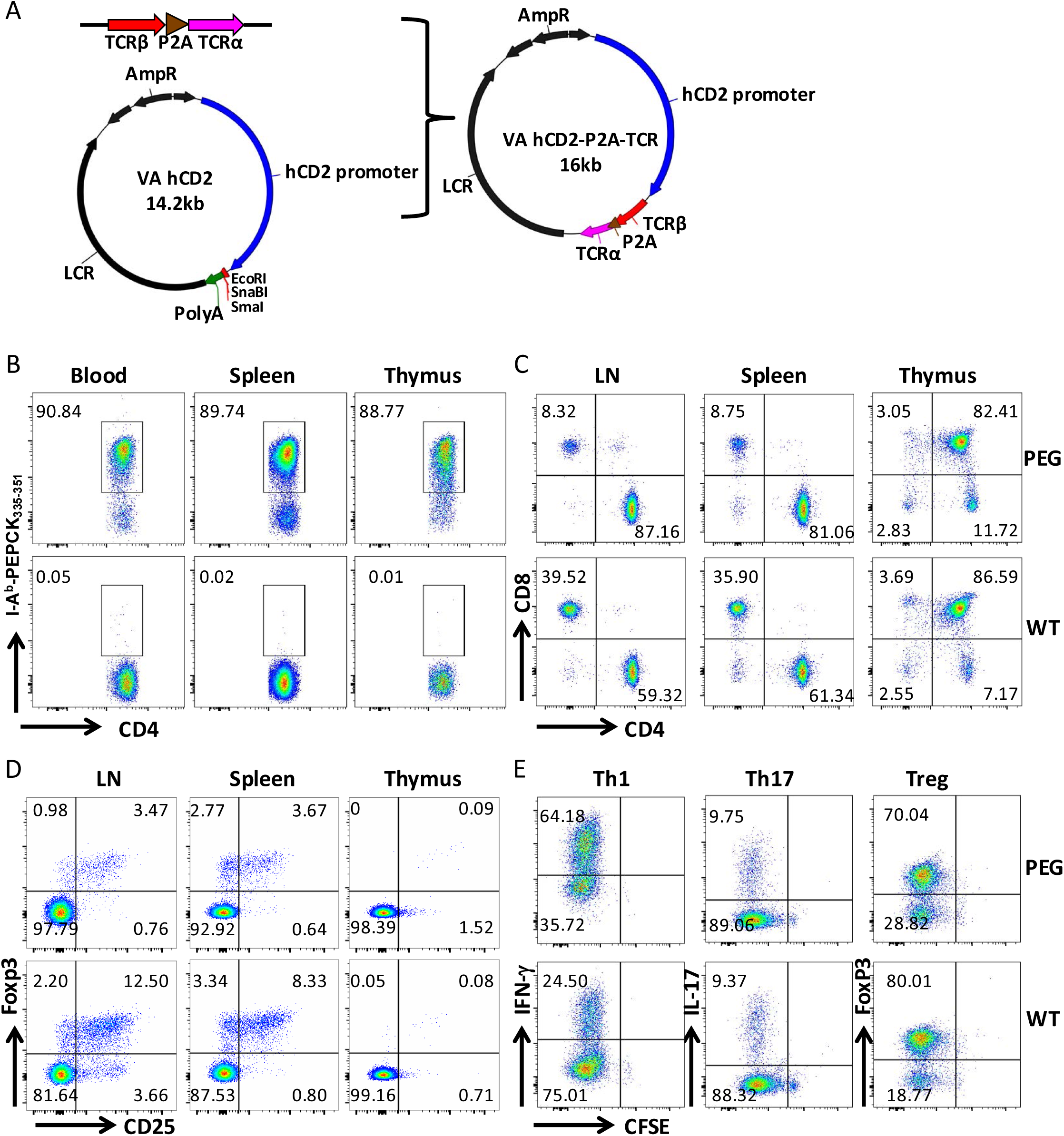
Generation and phenotyping of PEPCK-specific TCR Tg (PEG) mice. **(A)** Schematic protocol for constructing transgenic vector for generating PEG mice. PEPCK-specific TCR alpha and beta genes were linked with P2A linker and subcloned into the multiple cloning site of VA hCD2 vector under hCD2 promoter. **(B)** Transgenic TCR expression on CD4^+^ T cells in blood, spleen and thymus in WT and PEG mice. Six to eight weeks old WT and PEG mice were sacrificed and the expression of transgenic TCR on the CD4^+^ cells (ability to bind to the pMHC tetramer) in the blood, spleens and thymus was assessed by flow cytometry. The frequency of CD4^+^ and CD8^+^ T cells **(C)** and Foxp3^+^ and CD25^+^ cells **(D**) in the lymph nodes, spleens and thymus of PEG and their WT littermate controls were also assessed. Purified (from spleens) CD4+ PEG and WT littermate control cells were differentiated into Th1, Th17 and Treg cells in vitro and assessed by flow cytometry **(E).**

**Fig. S3.**
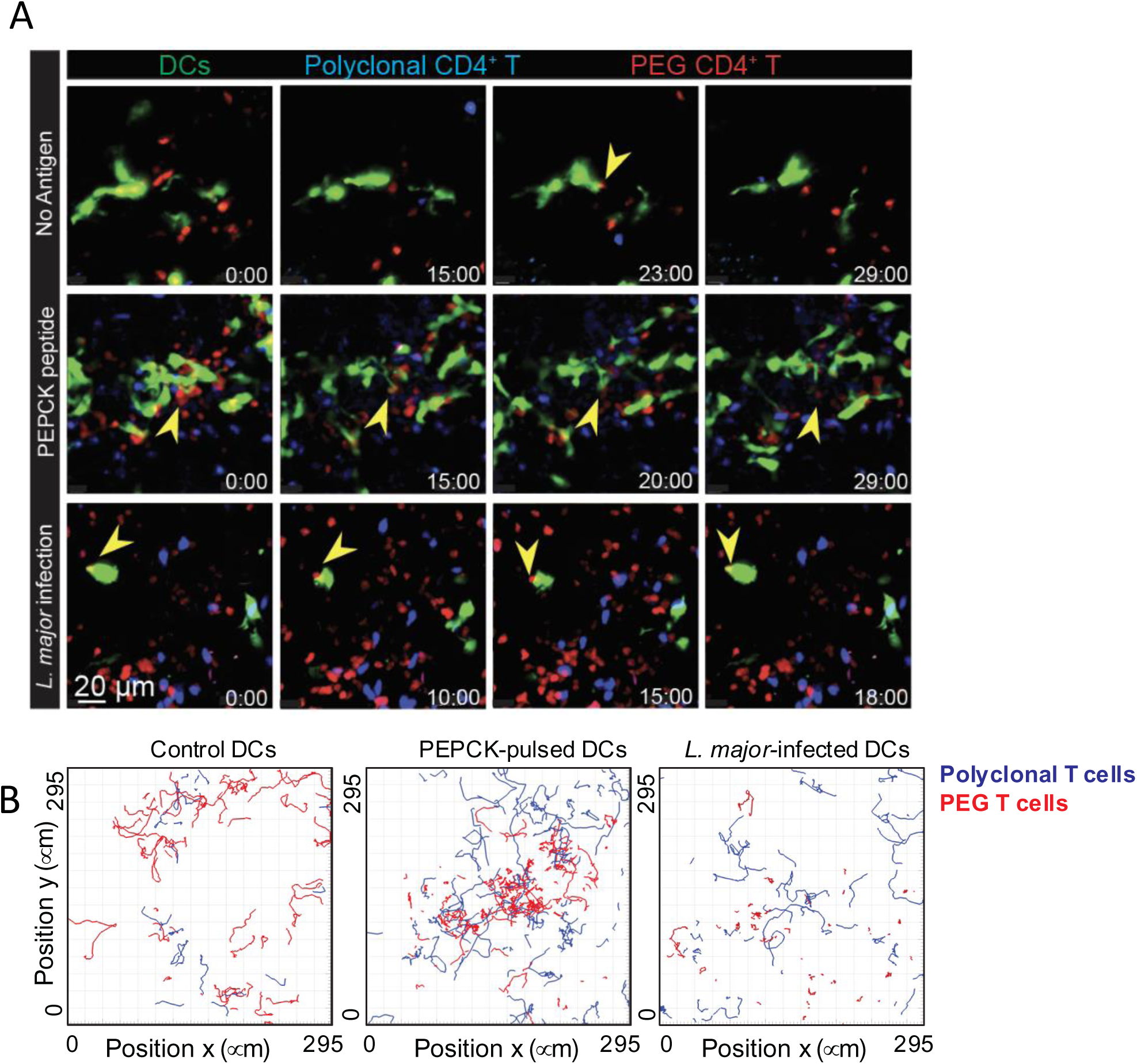
Migratory tracks of polyclonal (blue) and PEPCK-specific (red) CD4^+^ T cells *in vivo*. Bone marrow-derived dendritic cells were either pulsed with PEPCK_335-351_ peptides, infected with *L. major* or left untreated (uninfected), and then injected into the right hind footpads. After 12 hours, purified polyclonal or PEG TCR transgenic CD4^+^ T cells were labeled with Celltracker Blue or Celltracker Orange and injected into the mice. (A) Representative micrographs of the lymph nodes after adoptive transfer, yellow arrows represent T cell:DC contacts. Time stamps in min:sec, scale bar: 20 µm. (B) Migratory tracks of polyclonal and PEG populations during a 30-min recording.

**Fig. S4.**
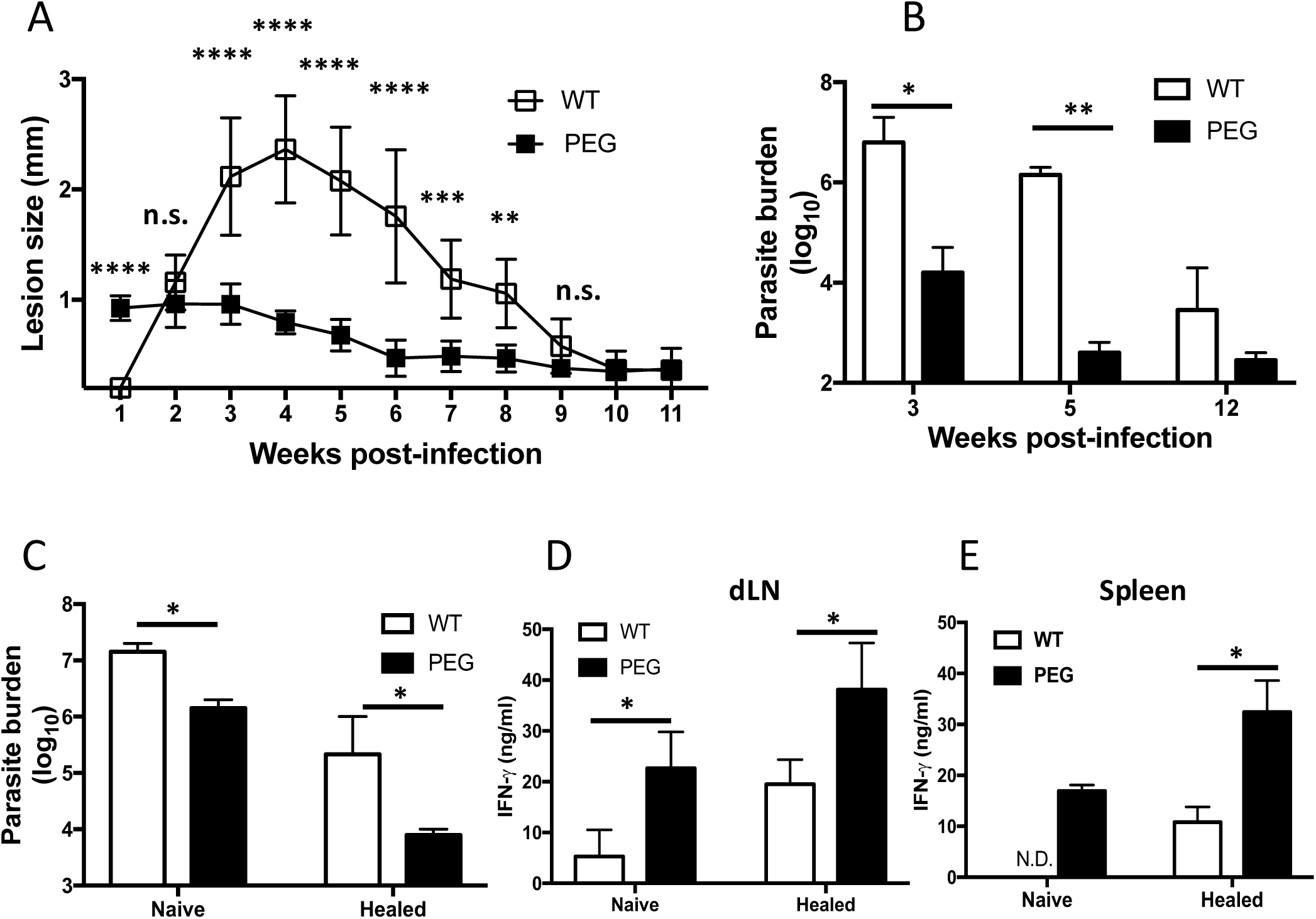
PEG mice are more resistant to *L. major* infection. PEG and their WT littermate control mice were infected with 1 × 10^6^ *L. major* in the right hind footpads and lesion size was monitored weekly with digital calipers (**A**). At the indicated times, mice were sacrificed, and parasite burden was measured by limiting dilution (**B**). After 12 weeks post-infection, naïve and healed PEG and WT littermate control mice were rechallenged with 1 × 10^6^ *L. major* in the left hind footpads. After 10 days, challenged mice were sacrificed and parasite burden was measured by limiting dilution (**C**). Cells from the draining lymph nodes (dLN, **D**) and spleens (**E**) were restimulated *in vitro* with SLA (50 μg/ml) for 3 days and the amount of IFN-γ in culture supernatant fluids of was measured by ELISA. *, p < 0.05; **, p < 0.01; ***, p < 0.001; ****, p < 0.0001; ns, not significant.

**Fig. S5.**
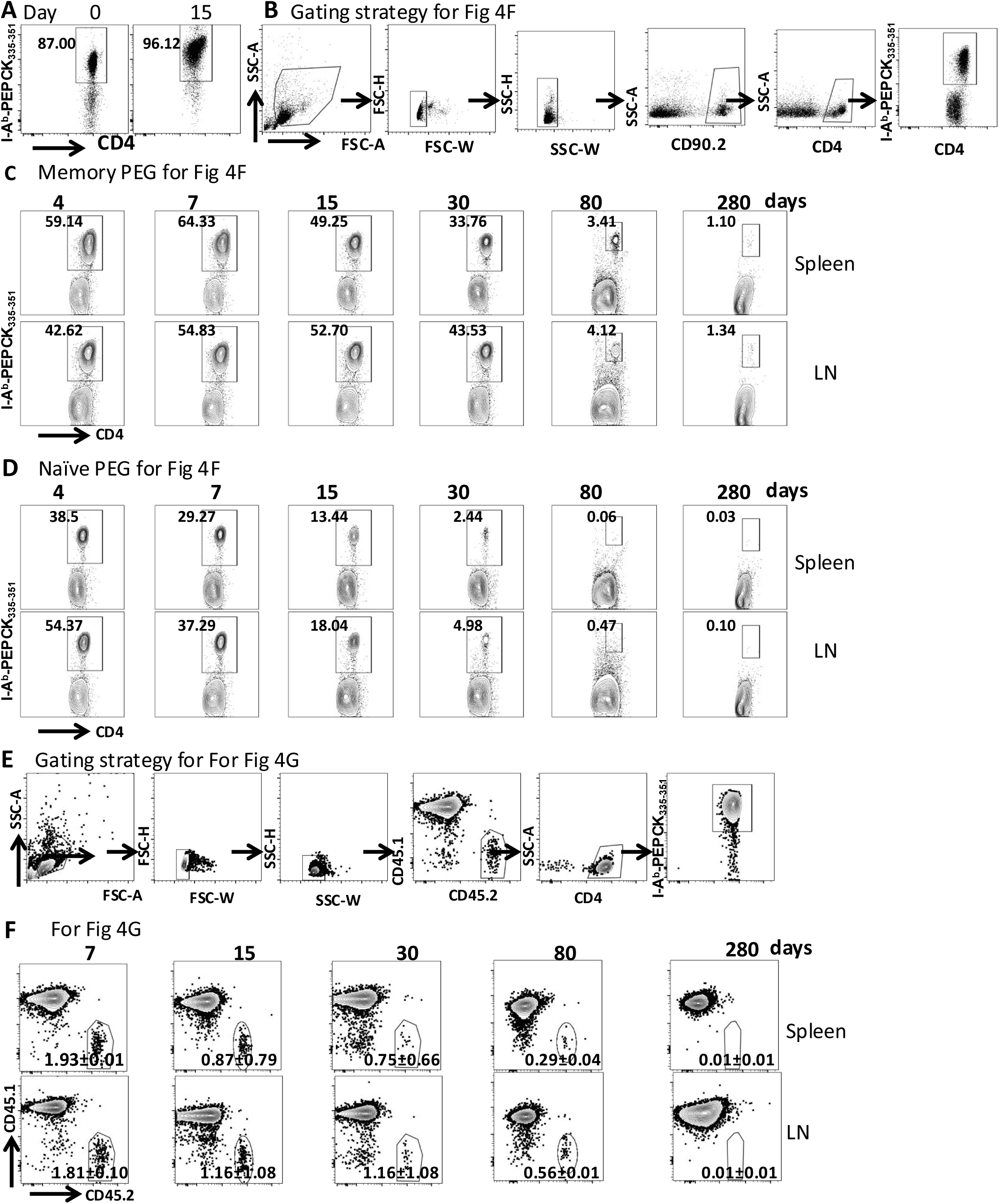
Supporting flow cytometry plots for Fig. 4. (**A**) transgenic TCR expression on naïve (Day 0) and *in vitro* generated memory (Day 15) PEG cells. (**B**) Gating strategy for the data presented as Fig. 4F. Flow cytometry detection of memory (**C**) and naïve (**D**) donor PEG cells in spleens and total peripheral lymph nodes (LN) in MHC II^-/-^ mice at indicated time points. (**E**) Gating strategy used to assess the numbers of PEG cells in congenic WT mice presented in Fig. 4G. (**F**) Flow cytometry detection of memory donor PEG cells in spleens and LNs in congenic WT mice at indicated the time points.

**Fig. S6.**
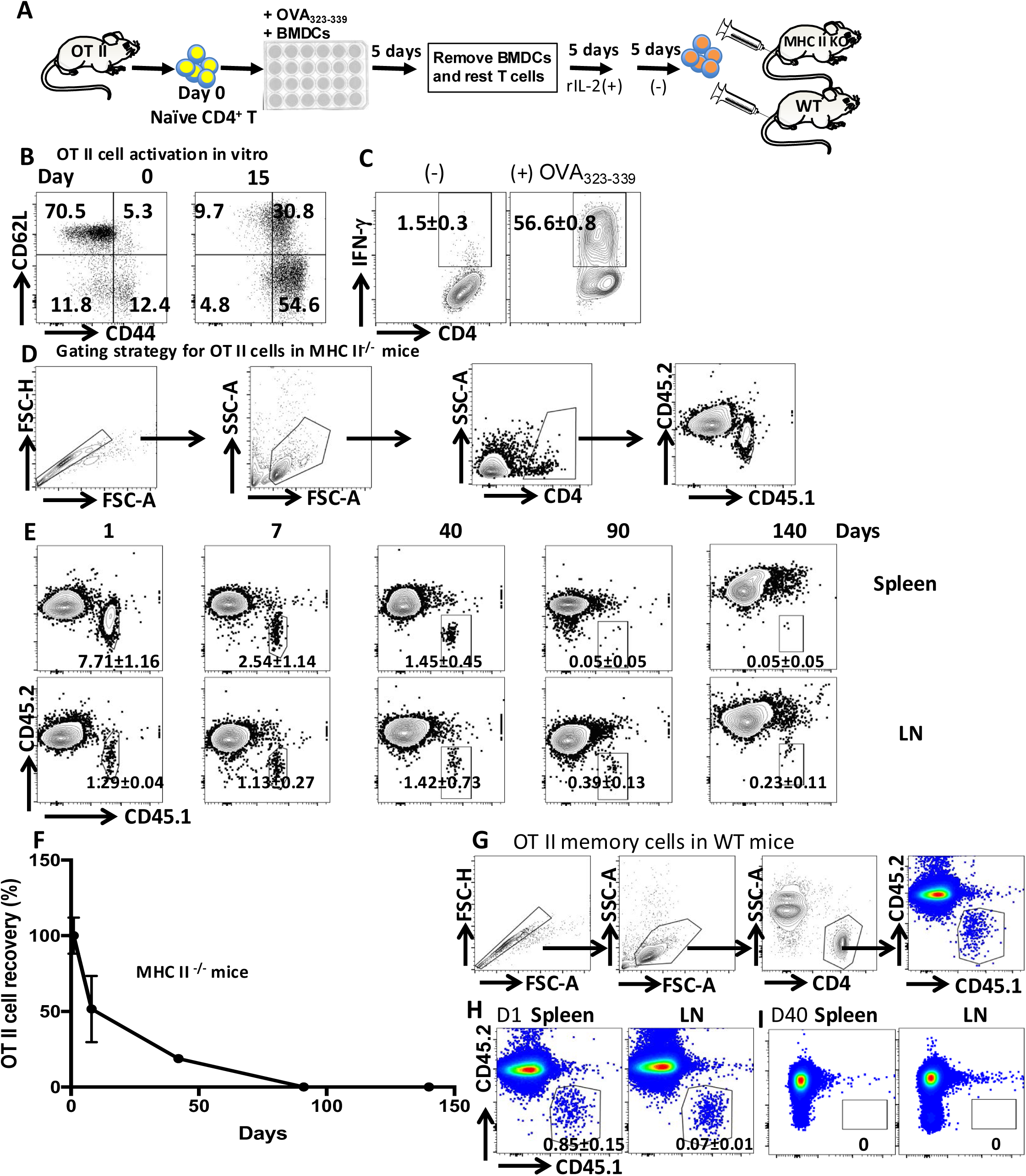
Memory OT II cells generated *in vitro* do not persist in MHC II^-/-^ and WT mice. (**A**) Schematic protocol for generating memory OT II cells *in vitro*. (**B**) Expression of memory markers on naïve (Day 0) and memory (Day 15) OT II cells. Memory OT II cells generated *in vitro* were stimulated overnight with splenic DCs pulsed with or without 5 μM OVA_323-339._ Thereafter, the cells were stimulated with BFA for additional 4-6 hr and IFN-γ production was assessed by flow cytometry (**C**). One million memory OT II cells generated *in vitro* were transferred into congenic MHC II^-/-^ mice and the indicated gating strategy was used to monitor OT II cells (**D**). At the indicated time points, the numbers of OT II cells in the recipient MHC II-/-mice were monitored by flow cytometry (**E** and **F**). One million memory OT II cells generated *in vitro* were transferred into congenic WT mice and their numbers were monitored at indicted time points using the depicted gating strategy (**G** and **H**).

**Fig. S7.**
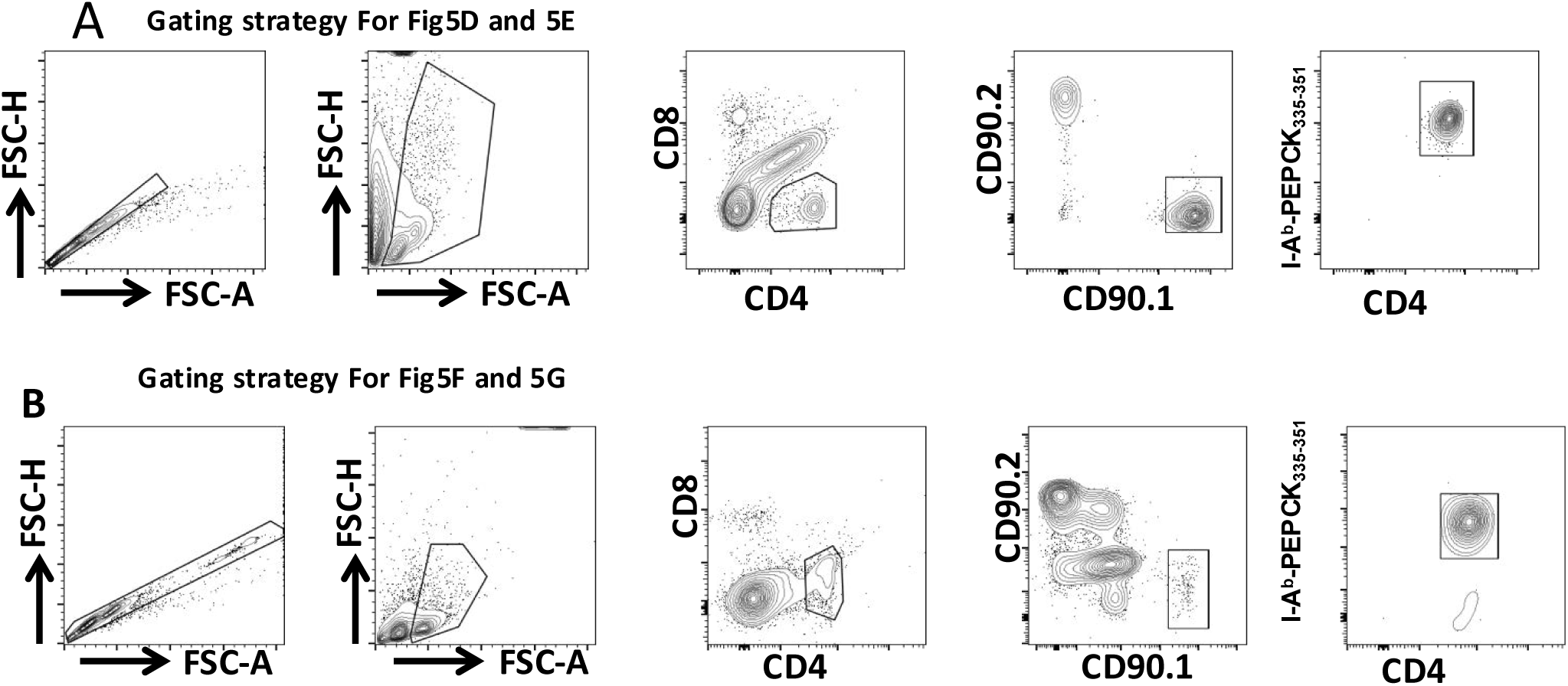
Gating strategies for Fig. 5. (**A**) Gating strategy for Fig. 5D and 5E. (**B**) Gating strategy for Fig. 5F and 5G.

**Fig. S8.**
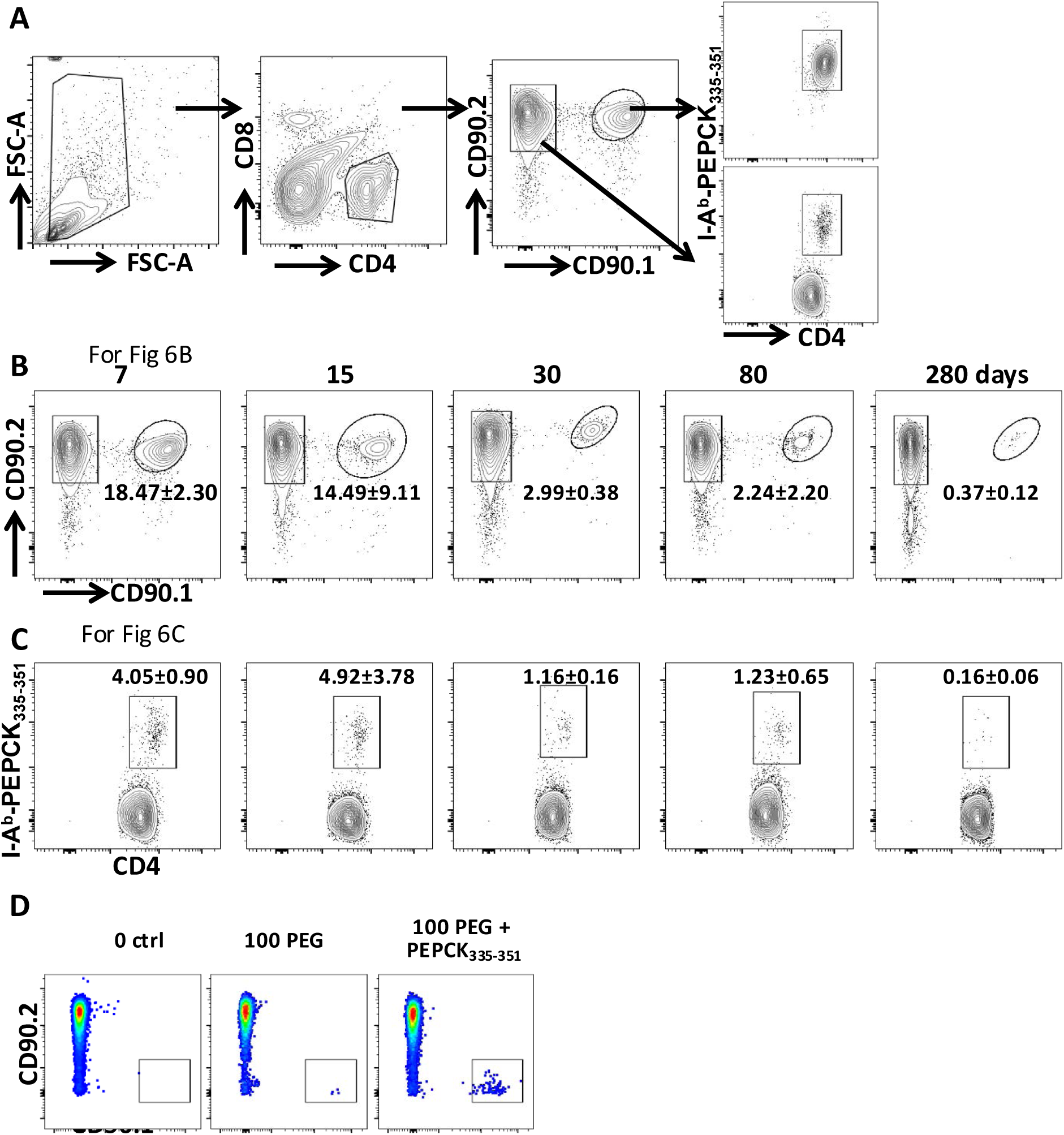
Supporting flow cytometry plots for Fig. 6. Ten thousand (10^4^) purified naïve PEG cells were transferred into congenic mice that were then injected with 5 nmol PEPCK_335-351_ and 25 μg CpG in the footpads the next day. (**A**) Gating strategy showing the assessment of donor (PEG) and endogenous (congenic) PEPCK-specific CD4^+^ T cells in the recipient mice. At the indicated times, the percentages of donor PEG cells (**B**) and endogenous PEPCK-specific CD4^+^ T cells (**C**) were monitored using magnetic beads-based enrichment with I-A^b^-PEPCK_335-351_ tetramer and analyzed by flow cytometry. (**D**) Magnetic bead-based tetramer enrichment is highly sensitive. One hundred (100) PEG cells were transferred into congenic mice and the recovery of the transferred cells was assessed in recipient mice next day by magnetic bead tetramer enrichment technique using I-A^b^-PEPCK_335-351_ tetramer. Some recipient mice were injected with 5 nmol PEPCK_335-351_ and 25 μg CpG and PEG cell numbers were assessed at 7 days post peptide challenge. Naive mice that were not injected with PEG cell were used as negative controls.

**Fig. S9.**
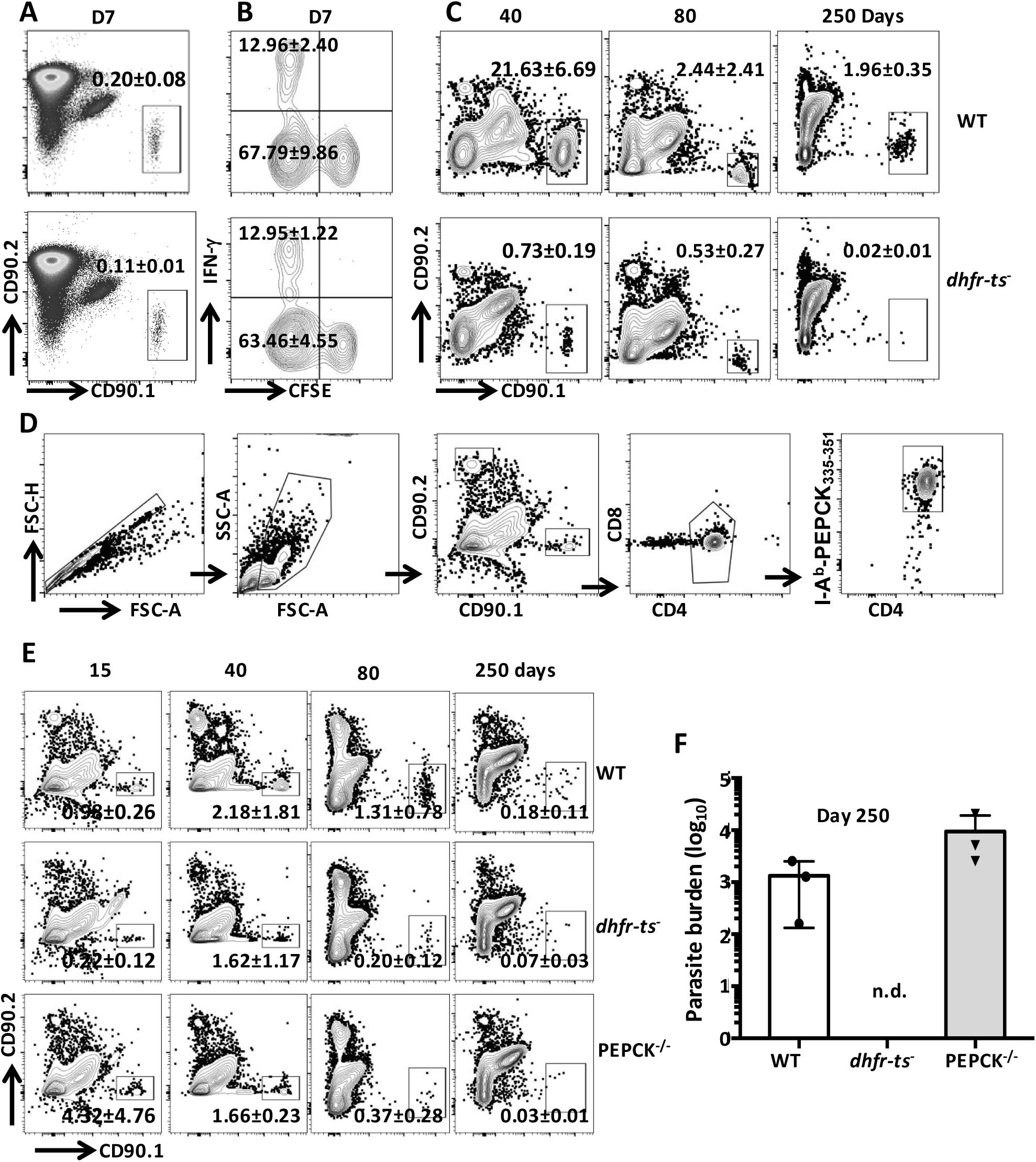
Supporting flow cytometry plots for Fig. 7. (**A-C**) Supporting flow plots for Fig. 7B. Ten thousand (10^4^) CFSE-labelled naïve PEG cells were transferred into congenic mice that were then infected with 10^6^ wild type (WT) or auxotrophic *dhfr-ts^-^ L. major* in the footpads the next day. (**A-B**) Early (day 7 post transfer) PEG cell responses to WT or *dhfr-ts^-^ L. major* were measured at 7 days post-infection. PEG cell numbers (**A**) and cell proliferation and IFN-γ production (**B**) were analyzed by flow cytometry. (**C**) Donor PEG cells were further monitored by flow cytometry at the indicated times. (**D-F**) Supporting flow plots for Fig. 7F. Ten thousand (10^4^) naïve PEG cells were transferred into congenic mice that were then injected with 5nmol PEPCK_335-351_ in presence of 25 μg CpG in right hind footpads the next day. After one week, the recipient mice were infected with 10^4^ WT, *dhfr-ts^-^* or PEPCK^-/-^ *L. major* in the left hind footpads and sacrificed at the indicated times to assess the numbers of PEG cells in the pooled peripheral lymph nodes cells and spleens. (**D**) Gating strategy for assessing the frequency of PEG cells for Fig. 7F. (**E**) The percentage of donor PEG cells in the recipient congenic mice was monitored by flow cytometry. (**F**) Parasite load was measured at 250 days post-infection by limiting dilution.

**Fig. S10.**
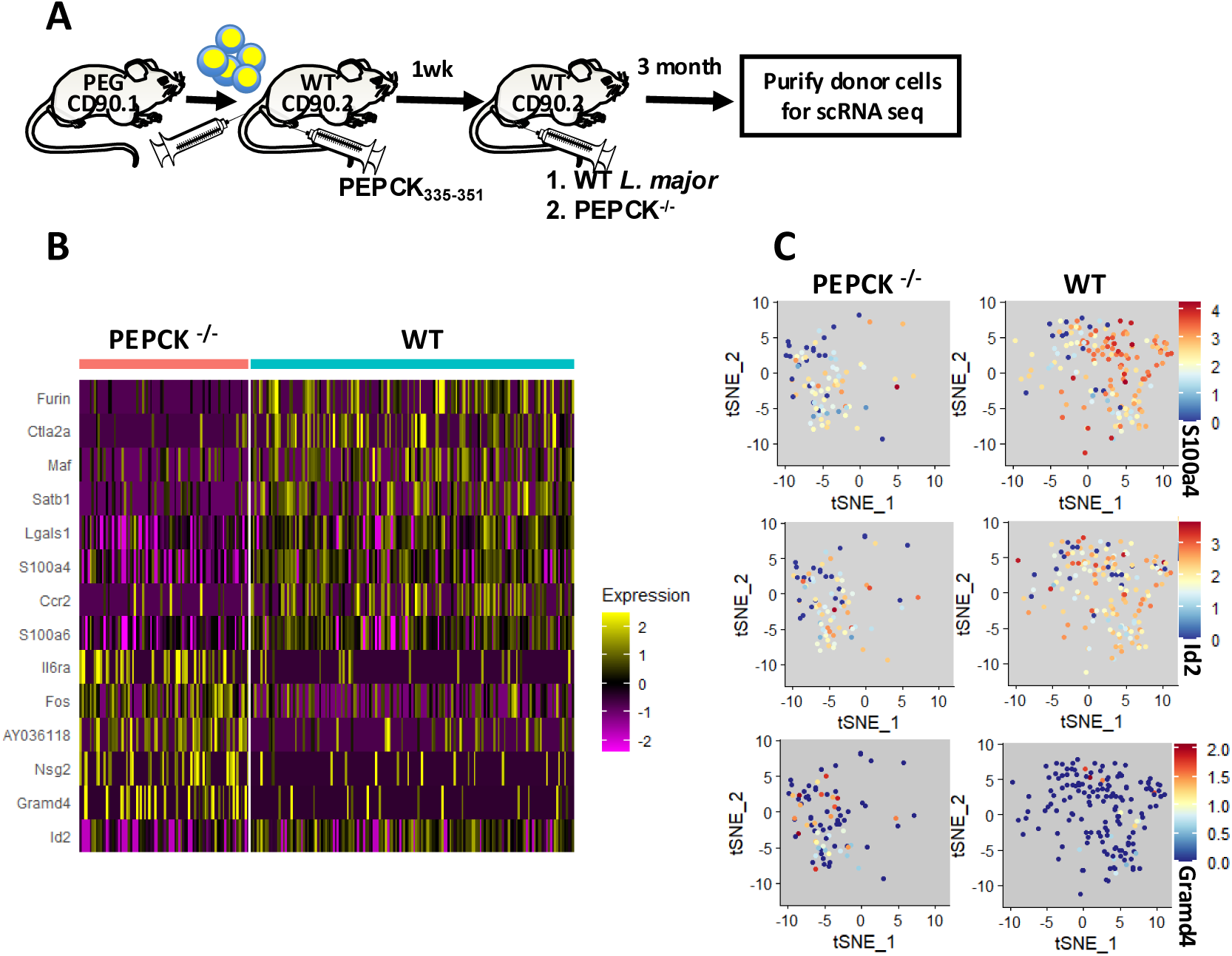
scRNA seq analysis of memory PEG CD4^+^ T cells in the presence or absence of their cognate antigen. **(A)** Schematic protocol. Ten thousand (10^4^) purified PEG cells were transferred into congenic mice that were then injected with 5 nmol PEPCK_335-351_ and 25 μg CpG in the footpads the next day. After 1 week, the mice were then infected with either 10^4^ WT (which persists and express PEPCK) or PEPCK^-/-^ (which persists but do not express PEPCK) *L. major* in the footpads. After 90 days, donor PEG cells were purified by cell sorting and gene expression profile of the cells were analyzed using 10X genomics pipeline. **(B)** Heatmap visualization of 14 selected differentially expressed genes. **(C)** Scatter plots of S100a4, Id2 and Gramd4 expression in memory PEG CD4^+^ T cells in absence (PEPCK^-/-^) or presence (WT) of their cognate antigen.

**Fig. S11.**
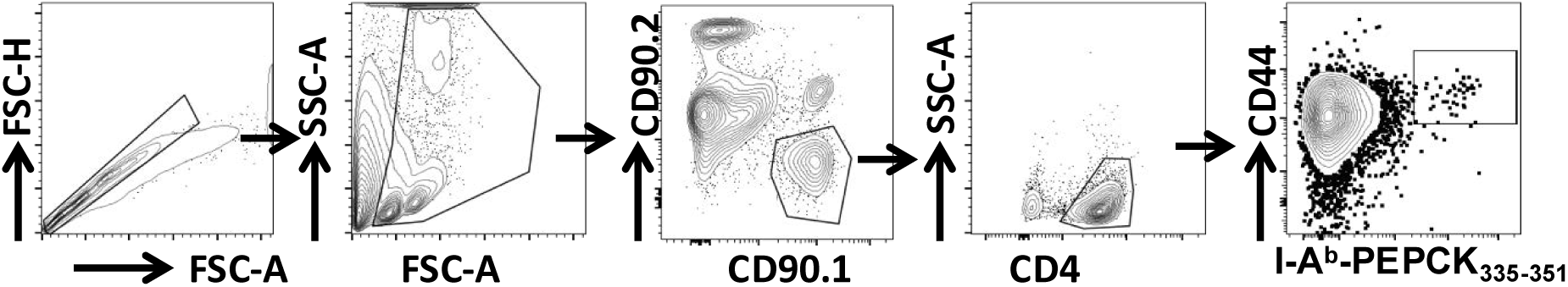
Gating strategy for measuring expansion of endogenous PEPCK-specific T cells for data presented in Fig. 8B-8C.

Movie S1. Cognate antigen recognition stabilizes DC-T cell conjugate formation in the lymph node.

## Acknowledgements

We acknowledge the NIH Tetramer Core Facility (contract HHSN272201300006C) for provision of I-A^b^–PEPCK_335–351_ tetramers. We thank A. Barazandeh and G. Akaluka for genotyping PEG mice.

## Funding

JU is supported by the Canadian Institutes for Health Research (CIHR Project Grant #178224). RZ is a recipient of a studentship from Research Manitoba.

## Author contributions

JU conceptualized the study. JU, ZM, TT and RZ designed the methodology. ZM, RZ, HH, ES, NI and SSO performed the experiments. ZM and RZ conducted the analysis. ZM and JU wrote the manuscript. JU, DL, HK, TT reviewed and edited the manuscript. JU acquired the funding and provided overall supervision.

## Competing interests

Authors declare that they have no competing interests.

## Data and materials availability

All data are available in the main text or supplementary materials. TCR transgenic mice generated in this study are available at University of Manitoba and can be shared upon completion of materials transfer agreements (MTAs).

## Notes

### Competing Interest Statement

The authors have declared no competing interest.

